# Parvalbumin Interneuron Activity Underlies Vulnerability to Post-Traumatic Stress Disorder in Autism

**DOI:** 10.1101/2021.01.18.427217

**Authors:** A. Shaam Al Abed, Tiarne V. Allen, Noorya Y. Ahmed, Azza Sellami, Yovina Sontani, Aline Marighetto, Aline Desmedt, Nathalie Dehorter

## Abstract

A rising concern in Autism Spectrum Disorder (ASD) is the heightened sensitivity to stress and trauma, the potential consequences of which have been overlooked, particularly upon the severity of the ASD traits. This study investigated the predisposition to, and impact of, Post-Traumatic Stress Disorder (PTSD) in ASD. We first demonstrated a reciprocal relationship between the two disorders and revealed that exposure to a mild stressful event induces PTSD-like memory in four mouse models of ASD. We also establish an unanticipated consequence of stress in this condition, showing that the formation of PTSD-like memory leads to the aggravation of the core traits associated with ASD. Such a susceptibility to developing PTSD-like memory in ASD stemmed from hyperactivation of the prefrontal cortex and altered fine-tuning of parvalbumin interneuron firing. We show that this traumatic memory can be treated by recontextualization, reducing the deleterious effects on the core symptoms of ASD. Overall, this study reveals multi-level neurobiological mechanisms that explain the increased vulnerability to develop PTSD in ASD. It provides a framework for future examination of the impact of PTSD-like memory in autism and offers new directions toward behavioral therapeutic interventions targeting traumatic memory in ASD.

## Introduction

Autism Spectrum Disorder (ASD) is a neurodevelopmental disorder caused by a combination of genetic and environmental factors that impair neuronal circuit function and lead to behavioural difficulties, such as altered social behaviour and repetitive movements ^1,2^. Beyond the core traits, ASD patients present with cognitive defects, including hyper-reactivity to sensory stimuli ^3^, abnormal fear conditioning ^4^, and altered declarative memory ^5^. Such a combination of impairments suggests a potential predisposition for developing post-traumatic stress disorder (PTSD), in which an extreme stress induces an altered memory of an event ^1,6^. In line with recent studies in humans positing a co-occurrence of these two disorders ^7–9^, ASD and PTSD display common behavioral features including impaired emotional regulation, cognitive rigidity, and fragmented autobiographical memory ^5^. Yet, the link between the two disorders remains poorly explored. This is critical to address as there is a pressing need for explaining the occurrence of traumatic stress in autism, but also for understanding the mechanisms underlying the unique perception of traumatic events in this condition ^7^.

As a crucial component of the fear memory circuitry ^10^, the medial prefrontal cortex (mPFC) controls downstream structures including the hippocampus and the amygdala to drive executive functions such as short-term memory and reasoning ^11,12^. The dysfunction of the mPFC has been shown in both ASD ^13^ and PTSD ^14^, suggesting that this structure represents a common node where cellular alterations may contribute to the emergence of these disorders. The mPFC contains the fast-spiking parvalbumin-expressing interneurons (PV-INs) that provide critical inhibitory control and maintain the excitation/inhibition balance in cortical circuits. These cells are required for adapted fear memorization ^11^ and normal sensory function ^15^. They play an essential role in stress-related disorders ^16^, as well as in ASD, where they drive aberrant cortical activity ^15,17,18^. Together, PV-INs represent cellular targets for normalizing functional connectivity and behavior in ASD, as their excitation has been shown to rescue social impairments in the Contactin-associated protein 2 knock out (*Cntnap2* KO) mouse model, which recapitulates the core symptoms of ASD ^18,19^. However, there is an urgent need to investigate the extent to which PV-INs are involved in the cognitive deficits and their role in tuning cortical activity in response to stress in ASD.

In this study, we aimed toimpactsl key cellular and molecular mechanisms involved in interneuron adaptation to stress that impact both medial prefrontal cortex activity and memory formation, and in turn contribute to the pathophysiology of ASD.

## Results

### Stress in memorization and autistic traits

No investigation has explored the connection between ASD and PTSD in mice models of ASD. A cardinal feature of PTSD is the development of a maladaptive memory of the traumatic event^20^. Here, we studied PTSD-like memory formation in the *Cntnap2* KO mouse model, using a contextual fear conditioning paradigm model of PTSD-like memory ^21,22^. This paradigm combines two auditory tones *unpaired* to two mild electric foot-shocks (**Fig. 1a; Day 1**), in which the tones do not predict the foot shocks. As such, the environment or “context” of the conditioning session becomes the only relevant predictor of the threat. We found that non-stressed control mice developed an adapted contextual fear memory, characterized by a low fear of the tone (*i.e.* tone test; **Fig. 1a, Day 2; Fig. 1b left panel**) combined with a strong fear of the conditioning context (*i.e.* context test; **Fig. 1a, Day 2; Fig. 1b right panel**). In contrast, submitting control mice to a 20 min restraint stress immediately after the conditioning led to the development of a maladaptive memory, consisting of a strong fear of the tone, and marginal fear of the conditioning context (**Fig 1b**). This abnormal memory profile mimics PTSD-related memory, by combining intrusive *sensory hypermnesia* for a salient, yet irrelevant element of the trauma (the tone), with a partial *amnesia* of the surrounding context ^1,6,23^. Strikingly, *Cntnap2* KO mice displayed the same memory profiles as control mice (**Fig. 1c**). Non-stressed *Cntnap2* KO mice were able to form adapted contextual memory, while stressed *Cntnap2* KO mice showed strong fear of the irrelevant tone, and low fear of the conditioning context. This demonstrates that, as control mice, *Cntnap2* KO mice develop PTSD-like memory after a stressful event.

**Fig. 1.**
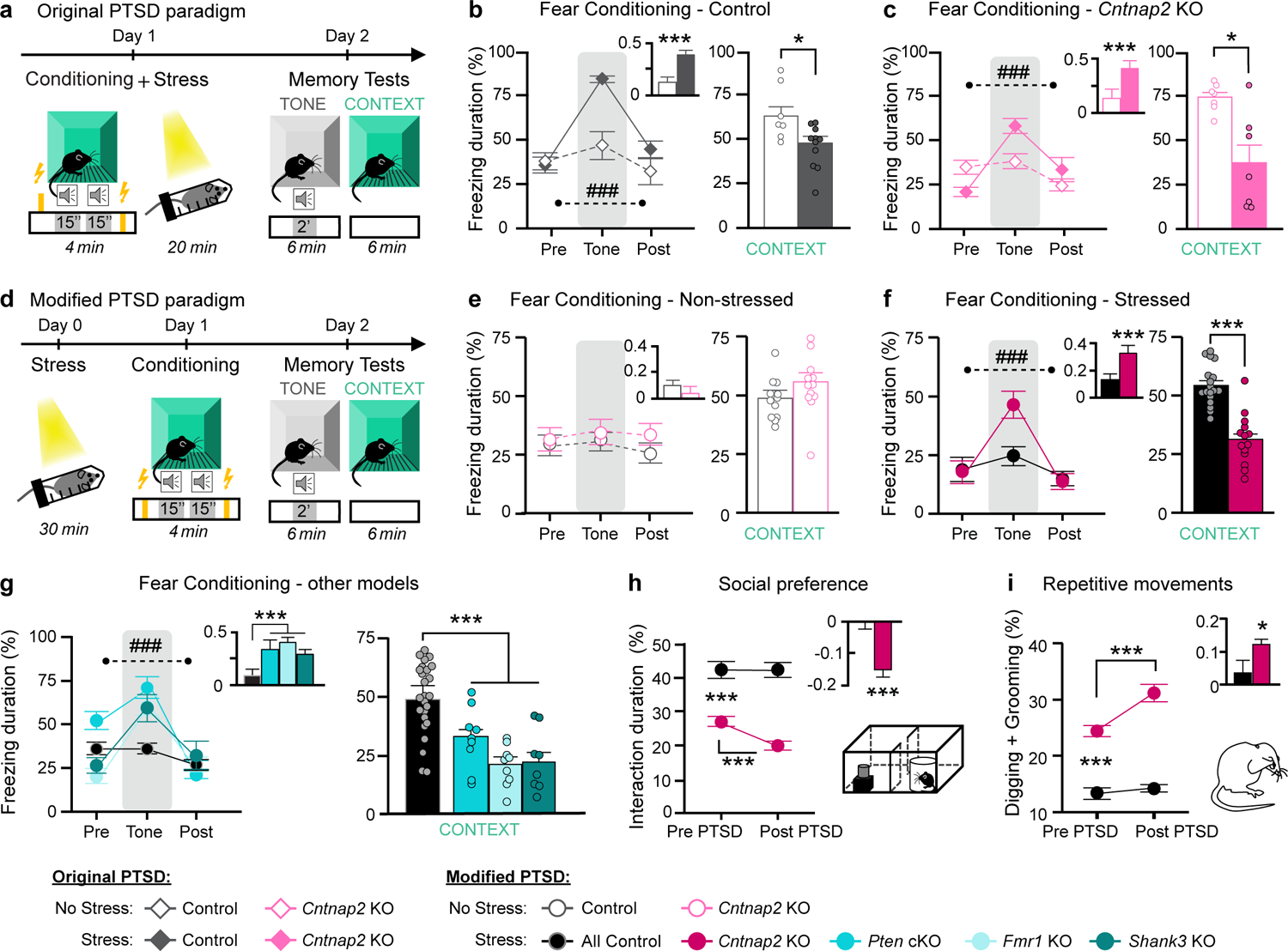
*Cntnap2* KO mice display traumatic memory profile after mild stress. **a**, Behavioral design of the original PTSD paradigm. **b**, Mean fear responses in control groups, measured as percentage of freezing duration before (pre), during (tone), and after (post) presentation of the tone (left; p<0.001; tone ratio (inset): p<0.001) and context tests (right; p<0.05); n=7 Non stressed Control (white fill) & 11 Stressed Control mice (grey fill). **c**, Mean fear responses in *Cntnap2* KO groups, measured as percentage of freezing duration before (pre), during (tone), and after (post) presentation of the tone (left; p<0.001; tone ratio (inset): p<0.001) and context tests (right; p<0.05); n=7 Non stressed *Cntnap2* KO (white fill) & 7 Stressed *Cntnap2* KO mice (pink fill). **d**, Behavioral design of the modified PTSD paradigm. **e**, Mean fear responses in non-stressed groups, measured as percentage of freezing duration before (pre), during (tone) and after (post) the tone (left; ns; tone ratio (inset): ns) and context tests (right; ns); n=18 Control (grey) & 20 *Cntnap2* KO mice (pink). **f**, Mean fear responses in stressed groups during tone test (left; p<0.001; tone ratio (inset): p<0.001) and context tests (right; p<0.001); n=32 Control (black) & 34 *Cntnap2* KO (magenta). **g**, Fear responses in stressed *Pten* control (n=7), *Pten* cKO (cyan; n=8), stressed *Fmr1* control (n=12), *Fmr1* KO (light blue; n=9) mice, stressed *Shank3* control (n=7), stressed *Shank3* KO (teal; n=8) mice (all controls *vs.* ASD models: tone test (left): p<0.0001; tone ratio (inset): p<0.001; context test (right): p<0.001). **h**, left; Social preference test before and after the modified PTSD paradigm: duration of interaction with an unfamiliar mouse in stressed control (black; n=11) and *Cntnap2* KO mice (Magenta; n=14; Control vs KO: p<0.001; Pre *vs.* Post PTSD in Control: ns and in KO: p<0.001; interaction ratio (inset): p<0.001). Bottom right: 2-chamber arena for social preference testing. **i**, Repetitive movements before and after the modified PTSD paradigm (digging and grooming; n=11 Control & 18 KO; Control vs KO: p<0.001; Pre-Post PTSD in Control: ns and in *Cntnap2* KO: p<0.001; ratio (inset): p<0.05). Data presented as mean ± SEM. ***: p<0.001; **: p<0.01; *: p<0.05 from 2-way ANOVA; #: interaction significance between freezing and genotype. See table for precise statistics.

It is well-known that ASD is characterized by difficulties coping with stress ^3^. We hence reasoned that PTSD-like memory could be triggered in *Cntnap2* KO mice with lower level of stress. Utilizing a modified version of the original PTSD protocol ^21,22^ we subjected mice to a 30 min restraint stress *24h before* conditioning (**Fig. 1d**). In this paradigm, both non-stressed control and *Cntnap2* KO mice displayed an adaptive contextual fear memory 24 hours post-conditioning (**Fig. 1d, Day 2; Fig. 1e**). In contrast, an independent group of stressed *Cntnap2* KO mice showed strong and specific fear response to the tone combined with low freezing to the conditioning context, whilst stressed control mice demonstrated normal contextual fear memory (**Fig. 1f**). This abnormal fear memory recapitulated all features of PTSD-like memory (Methods): partial fear generalization (*i.e.,* fear of a similar, yet non-identical, salient element of the traumatic event; **Fig. S1a**) and persisting across time (**Fig. S1b**). Furthermore, we found no difference in the reactivity to the electric foot-shock or the tone presentation between control and *Cntnap2* KO mice following stress and fear conditioning (**Fig. S1c**), indicating that PTSD-like memory did not stem from altered sensory processing ^3^. To confirm that this profile was a feature of the autistic condition and not due to specific features of the *Cntnap2* KO mouse model, we tested the protocol (*i.e*., restraint stress and unpaired fear conditioning) in three additional mouse models of ASD: The phosphatase and tensin homolog conditional knockout (*Pten* cKO), the *Shank* 3 KO, and the Fragile-X *Fmr1* KO mouse models of autism ^24,25^. Following this paradigm, *Pten* cKO, *Shank3* KO, and *Fmr1* KO mice displayed traumatic memory profile (**Fig. 1g**), overall proving a general vulnerability in autism to developing PTSD-like memory.

Given the similar behavioral features of both PTSD and ASD conditions (impaired emotional regulation, cognitive rigidity ^26^), we hypothesized that PTSD-like memory formation could exacerbate ASD-related traits. We therefore quantified social (**Fig. 1h; Fig. S2a-b**) and repetitive behaviors of the *Cntnap2* KO mice (**Fig. 1i; Fig. S2c-d**) following PTSD-like memory formation, compared to stressed control mice. We found that unlike fear conditioning alone or stress alone, PTSD-like memory formation, with both the classic (**Fig. S2e-g**) or the modified PTSD protocol aggravated the severity of both core autistic traits (**Fig. 1h-i**). This exacerbation was maintained for 3 weeks post-conditioning (**Fig. S2f, g**).

We next examined the malleability of the pathological memory developed in ASD conditions to determine whether it was possible to rescue PTSD-like memory in ASD mice. We used a behavior-based rehabilitation strategy called “recontextualization” ^22,23,27^, which has been shown to successfully normalize PTSD-like memory in control mice ^22^. We re-exposed the *Cntnap2* KO mice with PTSD-like memory to the original tone in the conditioning context with no foot shock (**Fig. 2a**). Whilst the stressed *Cntnap2* KO mice replicated the amnesia to the conditioning context, followed by a strong and high fear response to the tone during the recontextualization session (**Fig. 2b**), 24h later they exhibited normal, contextualized fear memory (**Fig. 2c**). Importantly, restoring normal memory improved social behavior and decreased repetitive movements to levels close to that of control mice (**Fig. 2d, e; Fig. S2h-j**). To note, we did not find any sexual dimorphism regarding the impact of stress in fear conditioning (**Fig. S3a, b**), or the aggravation of the core autistic traits with PTSD-like memory formation in the *Cntnap2* KO mice (**Fig. 3Sc, d**). Together, this data demonstrates that whilst PTSD-like memory exacerbates the core symptoms of ASD, it is possible to manipulate this pathological memory and subsequently alleviate ASD-related behavior. Therefore, it provides evidence for a reciprocal relationship between PTSD and ASD.

**Fig. 2.**
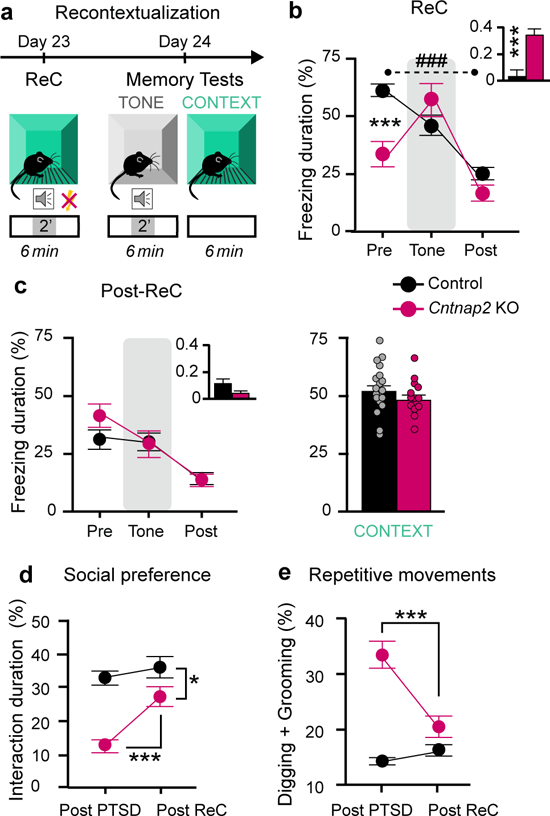
Recontextualization to normalize PTSD-like memory in ASD. **a**, Schematics of the recontextualization (ReC) session and subsequent memory tests. **b**, Fear responses during the Recontextualization (ReC) session (n= 19 control stress + FC and 14 *Cntnap2* KO stress + FC; Pre: p<0.001; #: interaction genotype x freezing: p<0.001; Tone ratio during ReC (inset): p<0.001). **c,** Fear responses post-ReC session, for the tone (left) and context (right) tests (n=19 Control and 14 *Cntnap2* KO mice; ns). **d**, Social preference before and after ReC protocol (n=10 Control & 12 KO; Post-PTSD *vs.* Post-ReC: ns in Control & p<0.001 in KO; Post ReC Control *vs.* KO p<0.05). **e**, Repetitive movements before and after ReC protocol (n=10 Control & 16 KO; Post-PTSD *vs.* Post-ReC: ns in Control & p<0.001 in KO). Data presented as mean ± SEM. ***: p<0.001; **: p<0.01; *: p<0.05 from 2-way ANOVA; #: interaction significance between freezing and genotype. See table for precise statistics.

### mPFC hyperactivation elicits PTSD-like memory

Alterations in the activity of the medial prefrontal cortex (mPFC) have been previously described in both PTSD ^14^ and ASD ^13,28^. Overall, decreased PFC functional activation to stressful and trauma-related cues, has been found in PTSD patients ^29^, while hyperactivity of the principal neurons has been described in ASD ^13^. To uncover the impact of PTSD-like memory formation in ASD, we analyzed neuronal activation induced by re-exposure to the conditioning context using the cell activity marker c-Fos ^21^. Whilst there was no difference in cFos-positive cell density at the basal level between control and *Cntnap2* KO mice (n= 4 mice/group; Data not shown), stressed *Cntnap2* KO mice presented more c-Fos positive cells than stressed control mice (**Fig. 3a, b**). This increase in mPFC cell activation was associated to alterations in downstream structures involved in the fear memory circuitry ^10^. We found a significant decrease in c-Fos-positive cells in the amygdala and hippocampus of stressed *Cntnap2* KO mice following re-exposure to the conditioning context, compared to control mice (**Fig. S4a, b**). Interestingly, limited cFos activation was detected in PV-INs, suggesting that the cFos increase observed after context test was specific to putative pyramidal cells (**Fig. 3a,b**).

**Fig. 3.**
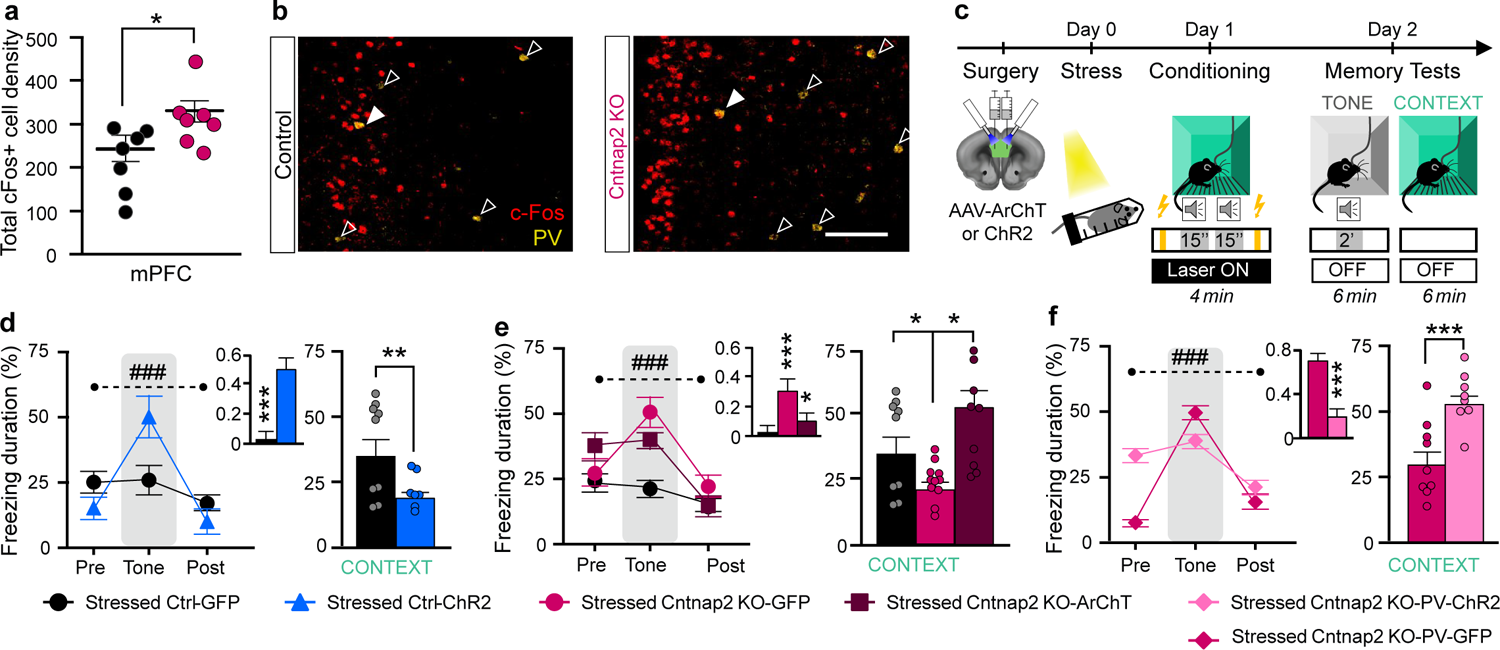
mPFC and PV-INs in the formation of PTSD-like memory. **a**, Total context recall-induced c-Fos hyperactivation in the medial pre-frontal cortex (mPFC; p<0.05); n= 7 Control & 7 *Cntnap2* KO mice. Proportion of PV+ cFos+ cells: 4.2 +/- 0.5% cFos+ PV+ in WT and 4.9 +/- 0.7 % in KO; 30 PV-INs out of 728 cFos+ cells in WT and 35 PV-INs out of 696 cFos+ cells in KO; p=0.482; 9.9 +/- 1.6% cFos+ PV-INs in WT and 7.8 +/-1.6 % in KO; 30 cFos+ out of 345 PV-INs cells in WT and 35 cFos+ out of 425 PV-INs cells in KO; p=0.352. **b,** Representative images of c-Fos density (red) in layer 2/3 of the mPFC (10x; scale: 100µm). **c,** *In vivo* optogenetics manipulation paradigm. **d,** Fear responses under photostimulation of the pyramidal neurons in ChR2-expressing control mice (n=7; blue) and GFP-infected control mice (n=9; black; tone test: p<0.001; tone ratio (inset): p<0.001; context test: p<0.01). **e,** Fear responses under photo-inactivation of the pyramidal neurons in ArChT-infected *Cntnap2* KO mice compared to GFP infected control and *Cntnap2* KO mice (*Cntnap2* KO ArChT: dark red, n=9; *Cntnap2* KO GFP: magenta, n=11, Control GFP: black, n=9; tone test: p<0.001; Pre-Tone, KO-GFP *vs.* ArChT: ns; tone ratio (inset): p<0.001, Control GFP *vs.* KO-GFP: p<0.001; KO-GFP *vs.* KO-ArChT: p<0.05; context test: control GFP *vs.* KO-GFP: p<0.05; KO-GFP *vs.* KO-ArChT: p<0.05). **f,** Fear responses under photostimulation of the Parvalbumin Interneurons (PV-INs) in ChR2-PV-Cre:*Cntnap2* KO mice compared to GFP-PV-Cre:*Cntnap2* KO mice (ChR2: light pink; n=9; GFP: magenta; n=9; tone test: p<0.001; tone ratio (inset): p<0.001; context test: p<0.001). Data are presented as mean ± SEM. ***: p<0.001; **: p<0.01; *: p<0.05; #: interaction significance between freezing and genotype. See table for precise statistics.

To further characterize the causal relationship between mPFC activity and PTSD-like memory formation, we performed *in vivo* optogenetic modulation of the pyramidal cells of the mPFC during the modified fear conditioning protocol **(Fig. 3c, Fig. S4c, d**). We found that optogenetic stimulation of the prefrontal cortical pyramidal cells via channelrhodopsin (ChR2) activation in stressed *control* mice was sufficient to trigger traumatic memory formation (**Fig. 3d**). Similarly, optical stimulation of the mPFC pyramidal cells in non-stressed C*ntnap*2 KO mice induced PTSD-like memory (**Fig. S4e**). Conversely, optogenetic inhibition of the mPFC pyramidal cells via Archaerhodopsin (ArchT) activation in stressed *Cntnap2* KO mice during conditioning successfully prevented PTSD-like memory formation (**Fig. 3e**). Together, this data indicates that stress-induced mPFC hyperactivation in ASD underlies PTSD-like memory formation.

### Stress-induced PV-INs alterations

Since an alteration of the excitation/inhibition balance coming from parvalbumin interneuron (PV-INs) hypofunction has been well documented in ASD ^18^, we then wondered whether PV interneuron activity underpins the observed mPFC hyperactivation and subsequent PTSD-like memory formation. We hypothesized that increasing interneuron activity would prevent traumatic memory formation. We performed optogenetic stimulation of PV-INs via cre-dependent ChR2 activation during conditioning in stressed *Cntnap2* KO mice and found that both PTSD-like memory formation (**Fig. 3f, Fig. S4f-i**), and the subsequent aggravation of core autistic traits were prevented (**Fig. S4j, k**). We next explored the physiological mechanisms priming PV-INs to be vulnerable to stress, using *in vitro* electrophysiological recordings in *PV-Cre; Td-Tomato* (controls) and *Cntnap2*^-/-^; *PV-Cre; Td-Tomato* (KO) mice. As previously reported ^30,31^, PV cell number was unchanged between control and KO conditions (**Fig. S5a-c**) as were intrinsic properties, such as resting membrane potential and input resistance (**Table 1**). However, we observed substantial changes in PV-IN firing pattern after stress in control mice, compared to non-stressed control mice (**Fig. 4, grey and black**). As previously described ^32^, stress enhances PV-IN excitability in control condition (*i.e.* reduction in spike latency and increase in maximum firing frequency (**Fig. 4a-h; Fig. S5d-f**). Interestingly, our data shows that *Cntnap2* KO PV-IN firing resembles that of stressed control PV-INs (**Fig. 4b, black and light pink**). PV-INs in the stressed *Cntnap2* KO condition presented a shift in excitability, characterized by a large increase in latency to first spike (**Fig. 4c-e, Fig. S5d-e**) and a significant decrease in firing frequency at intermediate depolarizing steps, compared to non-stressed KO and control conditions (**Fig. 4f**). This suggests that PV-INs in the *Cntnap2* KO mice compute cortical information following stress differently from the control. These cells are likely to filter more efficiently a certain range of inputs, underlying a global weaker inhibitory power within the mPFC.

**Fig. 4.**
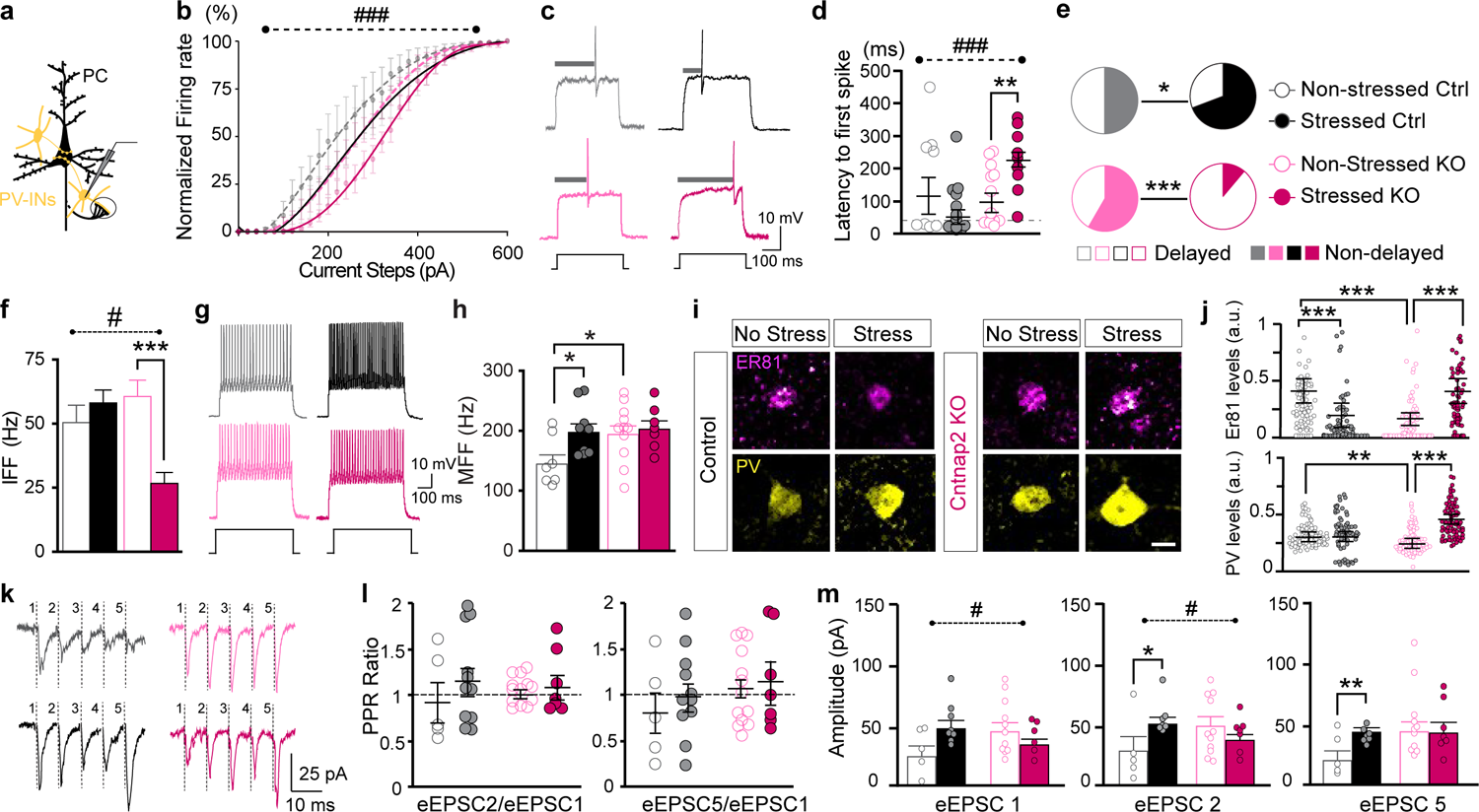
Effect of stress on PV-INs in controls and *Cntnap2* KO. **a**, Schematics of the *in vitro* electrophysiological recording of PV-INs in the mPFC; PC: Pyramidal Cell. **b,** Normalized PV-IN firing rate (% of maximum firing frequency) as a function of injected current steps (evolution of spike frequency x stress: p<0.001) for non-stressed controls (light grey; n=7), stressed controls (black; n=9), non-stressed *Cntnap2* KO (light pink; n=11), and stressed *Cntnap2* KO (dark pink; n=7) conditions. **c,** Representative traces of PV-IN firing at threshold potential. **d,** Latency to first spike (#: Interaction stress x genotype: p<0.001; stress in controls: ns; stress in *Cntnap2* KO: p<0.01). Dashed line represents limit (see Methods) between delayed *vs.* non-delayed cells. **e**, Proportion of delayed *vs.* non-delayed PV-INs in non-stressed and stressed conditions in control (Chi square test: p<0.05) and *Cntnap2* KO mice (p<0.001). **f,** Mean firing frequency at intermediate steps (IFF; 100-400 pA; #: stress x genotype: p<0.05; stress in controls: ns; stress in *Cntnap2* KO: p<0.001). **g,** Representative traces of PV-IN firing at maximum firing frequency (MFF, 600 pA). **h,** MFF at 600pA (stress in controls: p<0.05; non-stressed control *vs.* non-stressed KO: p<0.05). **i,** Representative illustrations of the expression of Er81 (Magenta) and PV (yellow) in each experimental group (scale: 10µm)**. j,** Mean Er81 (top) and PV (bottom) expression levels (a.u.: arbitrary unit; Er81: stress in controls: p<0.001; stress in *Cntnap2* KO: p<0.001; Genotype in non-stressed: p<0.001; PV: stress in controls: ns; stress in *Cntnap2* KO: p<0. 001; Genotype in non-stressed: p<0.01). **k,** Representative traces of 40 Hz paired-pulse ratios (PPR). **l,** PPR ratio between 1^st^ and 2^nd^ evoked excitatory post synaptic currents (eEPSC, left) and 1^st^ and 5^th^ EPSCs (right); n=5 no stress control; n=8 stressed control; n=11 *Cntnap2* KO no stress; n=7 *Cntnap2* KO stress; ns for both. **m,** Mean eEPSC amplitude (pA); #: interaction stress x genotype: p<0.05, p<0.05 & ns for EPSC1, EPSC2 & EPSC5, respectively; stress in control: ns, p<0.05 & p< 0.01; stress in *Cntnap2* KO: ns, ns, ns. Data are presented as mean ± SEM. ***: p<0.001; **: p<0.01; *: p<0.05. from 2-way ANOVA; see table for precise statistics.

**Table 1.** Electrophysiological properties of the parvalbumin interneurons of the prefrontal cortex in non-stressed and stressed control and Cntnap2 KO mice. Non-stressed PV-Cre; td-Tomato: n=7 cells, 4 mice; Stressed PV-Cre; td-Tomato: n=9 cells, 4 mice; Non-stressed *Cntnap2*^-/-^; PV-Cre; td-Tomato: n=10 cells, 5 mice; Stressed *Cntnap2*^-/-^; PV-Cre; td-Tomato: n=8 cells, 4 mice. 2-way ANOVA: * p<0.05

|  | Control |  | *Cntnap2* KO |  | p-values |
| --- | --- | --- | --- | --- | --- |
|  | Non-stressed | Stressed | Non-stressed | Stressed |  |
| **Vrest (mV)** | -65.63±1.81 | -65.89±1.47 | -71.96± 1.10 | -66.91±1.69 | Stress x Genotype: p=0.1201; Non-stress Ctrl vs. non-Stressed *Cntnap2* KO: p=0.0151 |
| **Rm (MΩ)** | 152.78±2.49 | 156.32±1.70 | 152.67±2.10 | 154.80 ±1.09 | Stress x Genotype: p=0.4413; Non-stress Ctrl vs. non-Stressed *Cntnap2* KO: p=0.7256; Stress in Controls: p=0.0724; Stress in *Cntnap2* KO: p=0.5928 |
| **V threshold (mV)** | -34.42±1.66 | -33.38±1.61 | -33.20±3.85 | -34.86±1.61 | Stress x Genotype: p=0.4929; Non-stress Ctrl vs. non-Stressed *Cntnap2* KO: p=0.6342; Stress in Controls: p=0.6323; Stress in *Cntnap2* KO: p=0.6253 |
| **AP amplitude (mV)** | 39.50±3.550 | 34.75±1.767 | 33.80± 2.390 | 27.01± 4.718 | Stress x Genotype: p=0.8904; Non-stress Ctrl vs. non-Stressed *Cntnap2* KO: p=0.1560; Stress in Controls: p=0.2145; Stress in *Cntnap2* KO: p=0.2579 |
| **AP rise time 10-90% (ms)** | 0.50 ± 0.07 | 0.61 ± 0.18 | 0.37 ± 0.05 | 0.34 ± 0.02 | Stress x Genotype: p=0.5415; Non-stress Ctrl vs. non-Stressed *Cntnap2* KO: p=0.1841; Stress in Controls: p=0.6258; Stress in *Cntnap2* KO: p=0.5111 |
| **AP half-width (ms)** | 0.79 ± 0.11 | 0.64 ± 0.04 | 0.61 ± 0.04 | 0.60 ± 0.03 | Stress x Genotype: p=0.4771; Non-stress Ctrl vs. non-Stressed *Cntnap2* KO: p=0.1793; Stress in Controls: p=0.1596; Stress in *Cntnap2* KO: p=0.4166 |
| **AP Decay 90-10% (ms)** | 0.58 ± 0.07 | 0.52 ± 0.05 | 0.50 ± 0.05 | 0.51 ± 0.05 | Stress x Genotype: p=0.9048; Non-stress Ctrl vs. non-Stressed *Cntnap2* KO: p=0.7658; Stress in Controls: p=0.5185; Stress in *Cntnap2* KO: p=0.4488 |
| **AHP peak (mV)** | -13.09 ± 1.33 | -12.08 ± 1.24 | -14.30 ± 1.19 | -14.55 ± 2.35 | Stress x Genotype: p=0.4563; Non-stress Ctrl vs. non-Stressed *Cntnap2* KO: p=0.4949; Stress in Controls: p=0.6080; Stress in *Cntnap2* KO: p=0.5948 |

These cellular alterations were associated with changes in the expression of activity-dependent proteins that fine-tune PV-IN firing, *i.e.* the calcium/calmodulin-dependent Etv1/Er81 transcription factor, which regulates PV-IN spike latency at near threshold potential ^33^, and the calcium-binding protein parvalbumin (PV; ^34^), which expression levels depend on cell activity ^35^. In line with the shifted firing latency (^33^; **Fig. 4c-e**), we found decreased Etv1/Er81 levels in control PV-INs upon stress (**Fig. 4i, j top panel**), but increased Etv1/Er81 expression in the *Cntnap2* KO condition after stress, compared to non-stressed condition.

As previously reported ^36^, PV levels were significantly decreased between control and *Cntnap2* KO conditions (**Fig. 4i, j bottom panel; Fig. S5c**). Whilst no change was observed upon stress in control, PV expression significantly increased in the stressed *Cntnap2* KO mice, compared to the non-stressed mice (**Fig. 4i, j bottom panel; Fig. S5c**). At the synaptic level, stress did not affect spontaneous inputs (**Fig. S5g, h**) and release probability onto PV-INs (**Fig. 4k, l; Fig. S5i, j**) in control and *Cntnap2* KO conditions. However, the amplitude of the evoked inputs afferent to PV-INs was increased upon stress in control (**Fig. 4m, black**). This was not the case for the *Cntnap2* KO condition (**Fig. 4m, pink**), suggesting that stress elicits differential regulation of the evoked inputs afferent to PV-INs. Overall, this data indicates that PV-INs in the *Cntnap2* KO mice display similar properties to stressed PV-INs in the control condition, and also present with an altered adaptation to stress that is likely to underlie a deficit in inhibition in the mPFC.

## Materials and methods

### Mice

3 to 5-month-old naive mice were individually housed in standard Makrolon cages, in a temperature- and humidity-controlled room under a 12h light/dark cycle (lights on at 07:00), with *ad libitum* access to food and water. We used both males and females to minimize the number of mice produced in this study. WT and *Cntnap2*^-/-^ KO mice (Jackson ID: #017482(Peñagarikano et al., 2011)) were non-littermates, derived from breeders of the same colony and used for behavior and histological analyses. *Shank 3* KO and control were kindly provided by Dr Mireille Montcouquiol and Dr Nathalie Sans (Neurocentre Magendie, Bordeaux University, France). *Pten* control (*Pten^F/F^*) and cKO mice (*Pten^F/F^; Emx-Cre/+*), and *Fmr1* control and KO mice were male littermates, kindly provided by Dr Andreas Frick (Neurocentre Magendie, Bordeaux University, France). For optogenetics manipulation of PV cells, PV-*Cre* mice (Jackson ID: #008069) were bred with *Cntnap2* ^-/-^ mice to obtain *Cntnap2* ^-/-^; PV-*Cre*/*Cre* (KO). For electrophysiological recordings, PV-*Cre* (Jackson ID: #008069); *Td-Tomato* mice (control) were bred with *Cntnap2* KO mice to obtain *Cntnap2* ^-/-^; PV-Cre/+; Td-Tomato/+ mice (KO).

All experiments took place during the light phase. We replicated the behavioral experiments in two to four different batches. We found no differences in the formation of PTSD-like memory in males and females (**Fig. S3**). Every effort was made to minimize the number of animals used and their suffering. All procedures were conducted in accordance with the European Directive for the care and use of laboratory animals (2010-63-EU) and the animals care guidelines issued by the animal experimental committee of Bordeaux University (CCEA50, agreement number A33-063-099; authorization N°21248), and from the Australian National University Animal Experimentation Ethics Committee (protocol numbers A2018/66, A2020/26, and A2021/43).

### Fear conditioning procedure

To assess the quality of the fear memory in the *Cntnap2* WT and KO mice, we used the only model currently available that allows the assessment of both memory components of PTSD in mice (*i.e.* emotional hypermnesia and contextual amnesia) (Kaouane et al., 2012).

#### Habituation (Day 0)

The day before fear conditioning, mice were individually placed for 4 min into a white squared chamber (30×15 cm, Imetronic, France) with an opaque PVC floor, in a brightness of 40 lux. The box was cleaned with 1% acetic acid before each trial. This pre-exposure allowed the mice to acclimate and become familiar with the chamber later used for the tone re-exposure test.

#### Acute mild stress

In the original protocol, a 20-min restraint stress was performed immediately after the conditioning. In the modified protocol, we performed a 30-min restraint stress under bright light (100 Lux) 24h before the conditioning session. Stressed mice were taken to a neutral room and placed into a perforated 50 mL Falcon® tube allowing air circulation. Non-stressed control mice were taken to the same room for 20 or 30 min but were kept in their home cage.

#### Conditioning (Day 1)

Acquisition of fear conditioning was performed in a different context, a transparent squared conditioning chamber (30×15 cm) in a brightness of 100 lux, given access to the different visual-spatial cues of the experimental room. The floor of the chamber consisted of 30 stainless-steel rods (5 mm diameter), spaced 5 mm apart and connected to the shock generator. The box was cleaned with 70% ethanol before each trial. All animals were trained with a tone-shock un-pairing procedure, meaning that the tone was non-predictive of the footshock occurrence. This training procedure, fully described in previous studies (Kaouane et al., 2012), promotes the processing of contextual cues in the foreground. Briefly, each animal was placed in the conditioning chamber for 4 min during which it received two tone cues (65 dB, 1 kHz, 15 s) and two foot-shocks (squared signal: 0.4 mA, 50 Hz, 1 s), following a pseudo-random distribution. Specifically, animals were placed in the conditioning chamber and receive a shock 100 s later, followed by a tone after a 30s interval. After a 20s delay, the same tone and shock spaced by a 30s interval were presented. Finally, after 20s, mice were returned to their home cage. As the tone was not paired to the footshock, mice selected the conditioning context (*i.e.* set of static background contextual cues and odor that constitutes the environment in which the conditioning takes place) and not the tone as the correct predictor of the shock (**Fig. 1a**, Day1).

#### Memory Tests (Day 2)

24 hours after conditioning, mice were submitted to two memory retention tests and continuously recorded for off-line second-by-second scoring of freezing by an observer blind to experimental groups. Mouse freezing behavior, defined as a lack of movement (except for respiratory-related movements), was used as an index of conditioned fear response (Fanselow, 1980). Mice were first submitted to the tone re-exposure test in the safe, familiar chamber during which three successive recording sessions of the behavioral responses were previously performed: one before (first two minutes), one during (next two minutes), and one after (two last minutes) tone presentation (**Fig. 1a**, Day 2). We determined the tone ratio (Insets in the figures) which corresponds to the conditioned response to the tone expressed by the percentage of freezing during tone presentation and compared to the levels of freezing expressed before and after tone presentation (repeated measures on 3 blocks of freezing). This tone ratio was calculated as follows: [% freezing during tone presentation – (% pre-tone period freezing + % post-tone period freezing)/2] / [% freezing during tone presentation + (% pre-tone period freezing + % post-tone period freezing)/2]. 2h later, mice were submitted to the context re-exposure test. They were placed for 6 min in the conditioning chamber. Freezing to the context was calculated as the percentage of the total time spent freezing during the successive three blocks of 2 min periods of the test. Optogenetic manipulation of memory formation was performed as previously described (Al Abed et al., 2020) and detailed in the optogenetics section of the methods.

### Assessment of the persistence of PTSD-like fear memory

To demonstrate that the PTSD-like memory was long-lasting, mice were again submitted to the two memory tests (tone-test and context-test spaced out of 2h, as in day 2), 20 days after fear conditioning (**Fig. S1b**). To assess generalization of fear, mice were exposed to 2 other tones, *i.e*, a new tone that is *similar* but not identical to the conditioning tone (3 kHz *vs.* 1kHz, respectively), and a white noise (*i.e.* highly *dissimilar* from the conditioning tone**; Fig. S1a**). We considered fear to be partially generalized when mice were presenting high levels of freezing to the new tone (3kHz).

**Recontextualization** was performed as previously described ^22^. Briefly, two days after long-term testing (Day 23), mice were re-exposed to the tone cue in the conditioning context, without electric shock (**Fig. 2ab**, Day 23). This protocol was also performed 3 days after conditioning, with similar results. The first 2 min of the recontextualization session (pre-tone) allowed us to assess the level of conditioned fear to the conditioning context alone, while the conditioned response to the tone was assessed during the next 2 min, both by the percentage of freezing during the tone presentation and by the tone ratio described above. 24h later, fear expression was assessed by re-exposing the mice to the regular (*i.e.* separated from each other) tone and context tests (same tests as in day 2).

### Assessment of the core symptoms of ASD

#### Repetitive behavior

Animals were placed in a cage with clean bedding and recorded for 5 min for off-line scoring. The behavior was defined by the percentage of time spent grooming for 2 sec minimum, as previously defined ^37^ and digging ^38^.

#### Social interactions

Following 2 min of habituation in an empty 2-chamber arena, we quantified the time spent close to a novel object or an unfamiliar, sex matched, control mouse for 5 min. The test mouse has access to an unfamiliar object and an unfamiliar mouse contained in a clear plastic box. Preference was quantified when the test mouse was located within a 5 cm-radius around the object/mouse. The results are expressed in percentage of duration of interactions with the mouse. Videos were analyzed off-line by an experimenter blinded to the experimental conditions. In addition to the main experimental group (stress + fear conditioning (**Fig. 1h, i**), we added control groups to ensure the specificity of the effect of traumatic memory on the severity of the core symptoms (*i.e.* mice were submitted to the stress or the fear conditioning only (**Supplementary** Fig. 2b-d**, f-g**)).

The exacerbation of core symptoms was studied according to the following timeline: Day 1: first assessment of ASD-like behaviors (i.e., social preference and digging/grooming). Day 2-4: fear conditioning. Day 5: second assessment of ASD-like behaviors. Day 6-7: Recontextualization. Day 8: third assessment of ASD-like behavior.

Long term maintenance of the ASD traits exacerbation was performed 3 weeks after the second assessment; no recontextualization session was performed in this case.

### Optogenetic manipulation of the mPFC

#### Surgery

Mice were injected bilaterally 4 weeks before behavioral experiments with an Adeno-Associated Virus (AAV) to inhibit (*AAV5-CaMKII*α*-ArchT-GFP*, UNC Vector Core) or activate glutamatergic neurons (*AAV5-CaMKII*α*-ChR2 (H134R)-EYFP*, UNC Vector Core) or parvalbumin interneurons (*AAV1-EF1a-DIO-hChR2(H134R)-eYFP*, UNC Vector Core). Control mice were injected with an AAV expressing GFP only (AAV5-CaMKIIα-GFP or *AAV1-CAG-flex-eGFP-WPRE.bGH,* UNC Vector Core). We used glass pipettes (tip diameter 25-35 µm) connected to a picospritzer (Parker Hannifin Corporation) into the mPFC (0.1 µl/site; AP +1.9 mm; L ± 0.35 mm; DV −1.3 mm). Mice were then implanted with bilateral optic fiber implants, 10 days before behavior (diameter: 200µm; numerical aperture: 0.39; flat tip; Thorlabs) directed to the mPFC (AP: +1.8 mm, L: ± 1.0 mm, DV: −1.3 mm, θ: 10°). Implants were fixed to the skull with Super-Bond dental cement (Sun Medical, Shiga, Japan). Mice were perfused after experiments to confirm correct placements of fibers (**Fig. S4c-e, g**). Viral injections targeted the L2/3 of the mPFC, and virtually all PV-INs in the prelimbic and infralimbic cortices were infected. For optogenetic manipulations, we used a LED (Plexon®) at 465nm with a large spectrum to allow the activation of both ArChT and ChR2. Light was continuously delivered to inactivate the mPFC, and was delivered at 5Hz (5ms ON,195ms OFF) for mPFC activation of the pyramidal cells (as previously described ^22^), and at 10Hz (10 ms ON, 10 ms OFF) for activation of the parvalbumin interneurons. Mice were submitted to the fear conditioning procedure described above and pyramidal cells of the mPFC were either inhibited or activated during the whole conditioning session. The next day, fear memory was tested as described above. To check the efficiency of the optogenetic activation of the PV-INs in the mPFC, we performed *in vitro electrophysiological* recordings of the pyramidal cells in current-clamp mode at −70mV (**Fig. S4h, i**).

### Immunohistochemistry

Animals were perfused transcardially 90 min after the context test on Day 2 for c-Fos analysis, and 60 min after the restraint stress for quantification of Etv1/Er81, and PV, with 0.01M phosphate buffered saline (PBS) to eliminate blood and extraneous material, followed by 4% paraformaldehyde (PFA). Brains were postfixed for 36h in PFA. Tissues were sectioned at 40 µm using a Leica 1000S vibratome and kept in a cryoprotective ethylene glycol solution at −20°C until processed for immunofluorescence. Sections were first washed and permeabilized in PBS-Triton 0.25%, then non-specific binding sites were blocked by immersing the tissue in 10% normal donkey serum, 2% bovine serum albumin in PBS-Triton 0.25% during 2h. Tissues were then stained using the primary antibodies overnight: mouse anti-c-Fos (1:1000; Santa Cruz), mouse anti-PV (Swan), rabbit anti-Etv1/Er81 (1:5000; kindly provided by Prof S. Arber, BioZentrum Switzerland). After 3x 15 min washes, we added anti-rabbit, anti-chicken, anti-mouse, Alexa 488 or 555 (1:200; Life Technologies) secondary antibodies for 2h. After 3x 15 min washes slices were stain during 10 min with DAPI (5µM; Sigma), mounted on Livingstone slides then covered with Mowiol (Sigma) and coverslip (Thermofisher). c-Fos, PV, and Er81 staining were imaged using an A1 Nikon confocal fluorescent microscope (20x objective). Stained sections of control and mutant mice were imaged during the same imaging session. Immunofluorescence signals were quantified using the ImageJ (FIJI) software with routine particle analysis procedures, to obtain nuclear masks, divided by the area to obtain cell density per mm^2^.

### In vitro Electrophysiology

Mice were deeply anaesthetized with isofluorane 60 min after the restraint stress, or from their homecage, and perfused with ice-cold oxygenated, modified artificial cerebrospinal fluid (ACSF) containing (in mM): 248 sucrose, 3 KCl, 0.5 CaCl2, 4 MgCl2, 1.25 NaH2PO4, 26 NaHCO3, and 1 glucose, saturated with 95% O2 and 5% CO2. The animals were then decapitated, the brain placed in icecold oxygenated modified ACSF and 300 μm coronal slices were cut using a Leica 1200S vibratome. Slices were then maintained at room temperature in ACSF containing (in mM): 124 NaCl, 3 KCl, 2 CaCl2, 1 MgCl2, 1.25 NaH2PO4, 26 NaHCO3 and 10 glucose saturated with 95% O2 and 5% CO2. For patch clamp recordings in whole-cell configuration, slices were transferred to a chamber and continuously superfused with ACSF at 32°C. We visualized interneurons located in layer 2-3 of the prefrontal cortex with infrared-differential interference optics through a 40x water-immersion objective. Microelectrodes (6–10 MΩ) were pulled from borosilicate glass (1.5 mm outer diameter x 0.86 inner diameter) using a vertical P10 puller (Narishige, Japan). We used a potassium-gluconate-based intracellular solution containing (in mM): 140 K-gluconate, 10 HEPES, 2 NaCl, 4 KCl, 4 ATP, and 0.4 GTP. Interneurons were kept under current-clamp configuration with an Axoclamp 200A (Axon Instruments) amplifier operating in a fast mode. Data were filtered on-line at 2 kHz, and acquired at a 20kHz sampling rate of using WinWCP5.5 software (Strathclyde University). Series resistance (Rs) was < 25 MΩ upon break-in and Δ Rs < 20% during the course of the experiment. To determine the excitability of the PV+ interneurons, we performed 500 ms depolarizing steps of 1 pA and used the first spike evoked by the minimum current needed to elicit an action potential applied from −70 mV. We considered a cell as “delayed” when the first elicited spike at threshold potential occurred more than 24.16 ms (*i.e.* median in non-stressed controls) after the beginning of the 1 pA depolarizing step (Dehorter et al., 2015).The following parameters were also measured: resting membrane potential (Vrest),membrane resistance, membrane capacitance (Cm), threshold potential for spikes (Vthreshold, defined as dV/dt = 10 mV/ms), rheobase, exponential fit of the slow ramp depolarization that remained just subthreshold during 500 ms current injections, AP amplitude, AP rise and AP duration, after-hyperpolarization (AHP) amplitude and AHP duration. We characterized the changes in membrane potential at near threshold (ΔVm), which corresponds to the activation of the delayed rectifier current (Golomb et al., 2007). We also performed 500 ms depolarizing steps of Δ25pA to determine the maximum firing frequency. Data analysis was performed off-line in EasyElectrophysiology (https://www.easyelectrophysiology.com/). Spontaneous excitatory postsynaptic currents (sEPSCs) were recorded in voltage-clamp mode at −70 mV. A stimulating electrode was placed in the cortex to activate cortical fibers and evoke PSCs in layer 2-3 cortical cells. Stimulus delivery was performed by ISO STIM 01D (NPI) and PSCs induced by a train of stimuli (10Hz, 20Hz and 40Hz, 5mA, 30 µs in deep layer V) were recorded in PV+ interneurons at −70 mV. For paired-pulse ratio (PPR), EPSC amplitude was measured on 5-10 averaged traces at each inter-pulse interval at the first (EPSC1), second (EPSC2) and fifth event (EPSC5).

### Statistics

Data are presented as mean ± SEM. Statistical analyses were performed using the StatView software for 2-way ANOVA, followed by Bonferroni *post-hoc* test or Student’s t test, when appropriate. Normality of the data was confirmed using the Kolmogorov–Smirnov test. Statistical significance between cell groups was performed using Chi square test (https://www.socscistatistics.com/). Statistical significance was considered at p<0.05. See **Table 2** (Main Figures) and **Table 3** (supporting information) for precise p-value and tests.

**Table 2.**
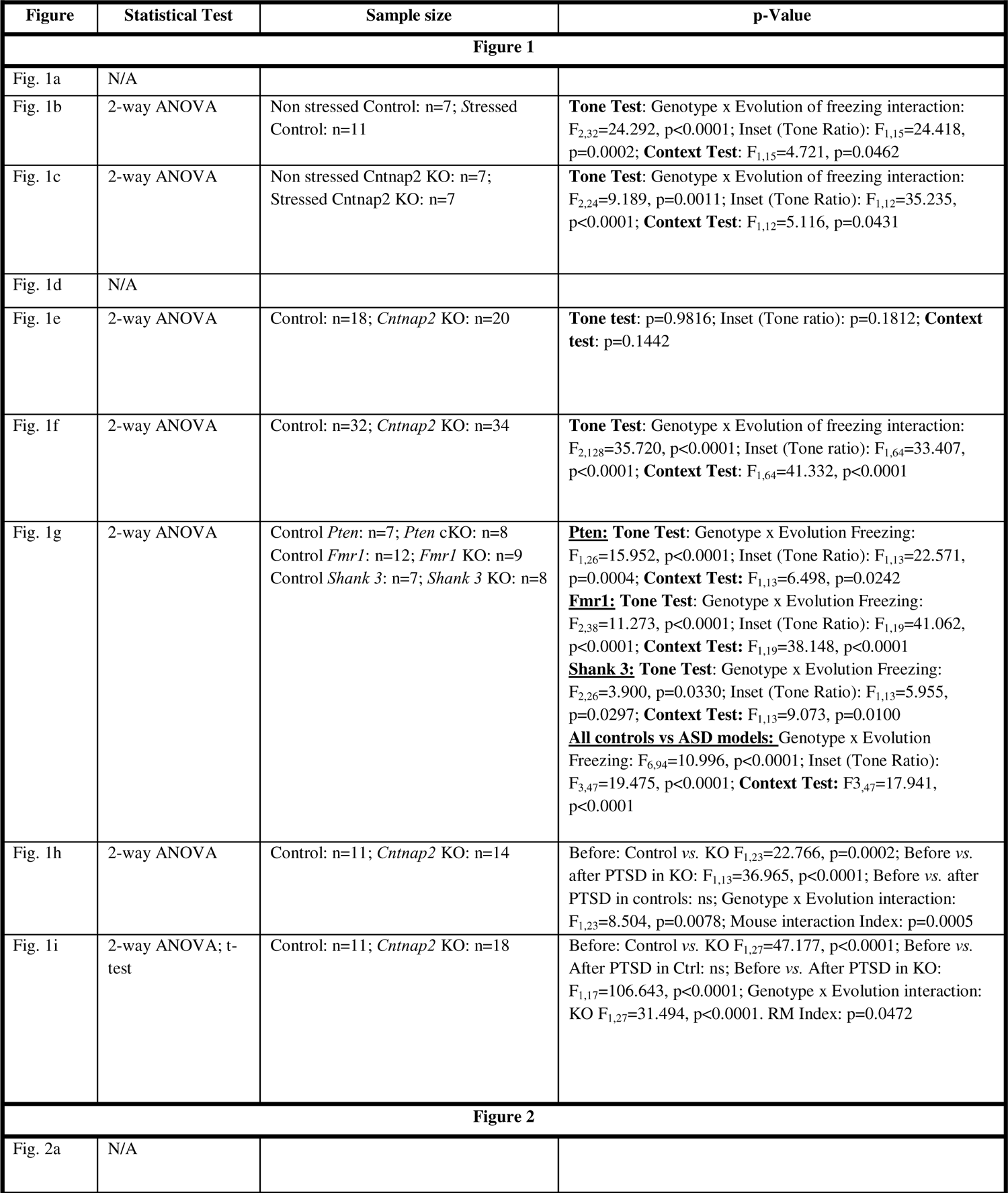
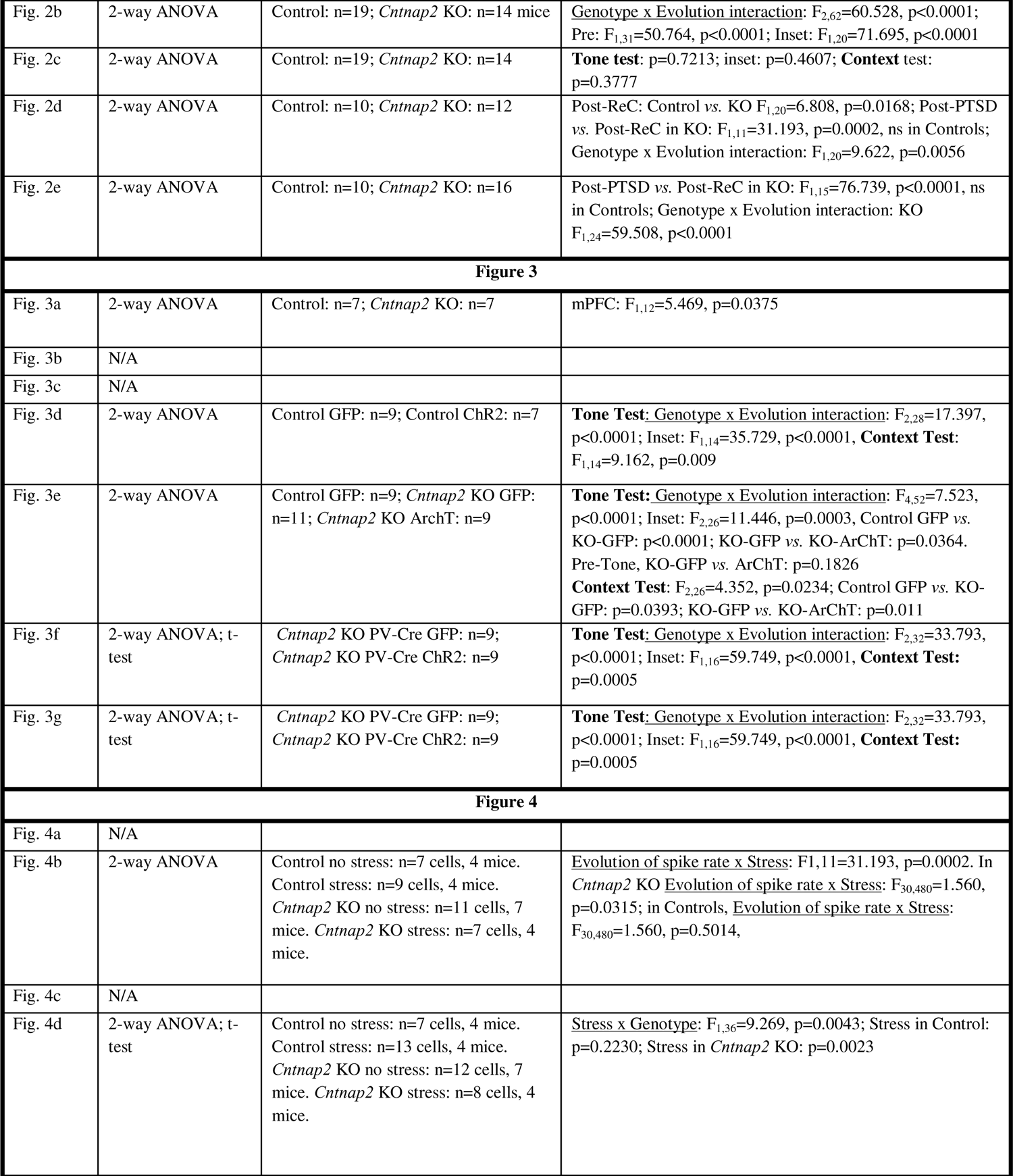
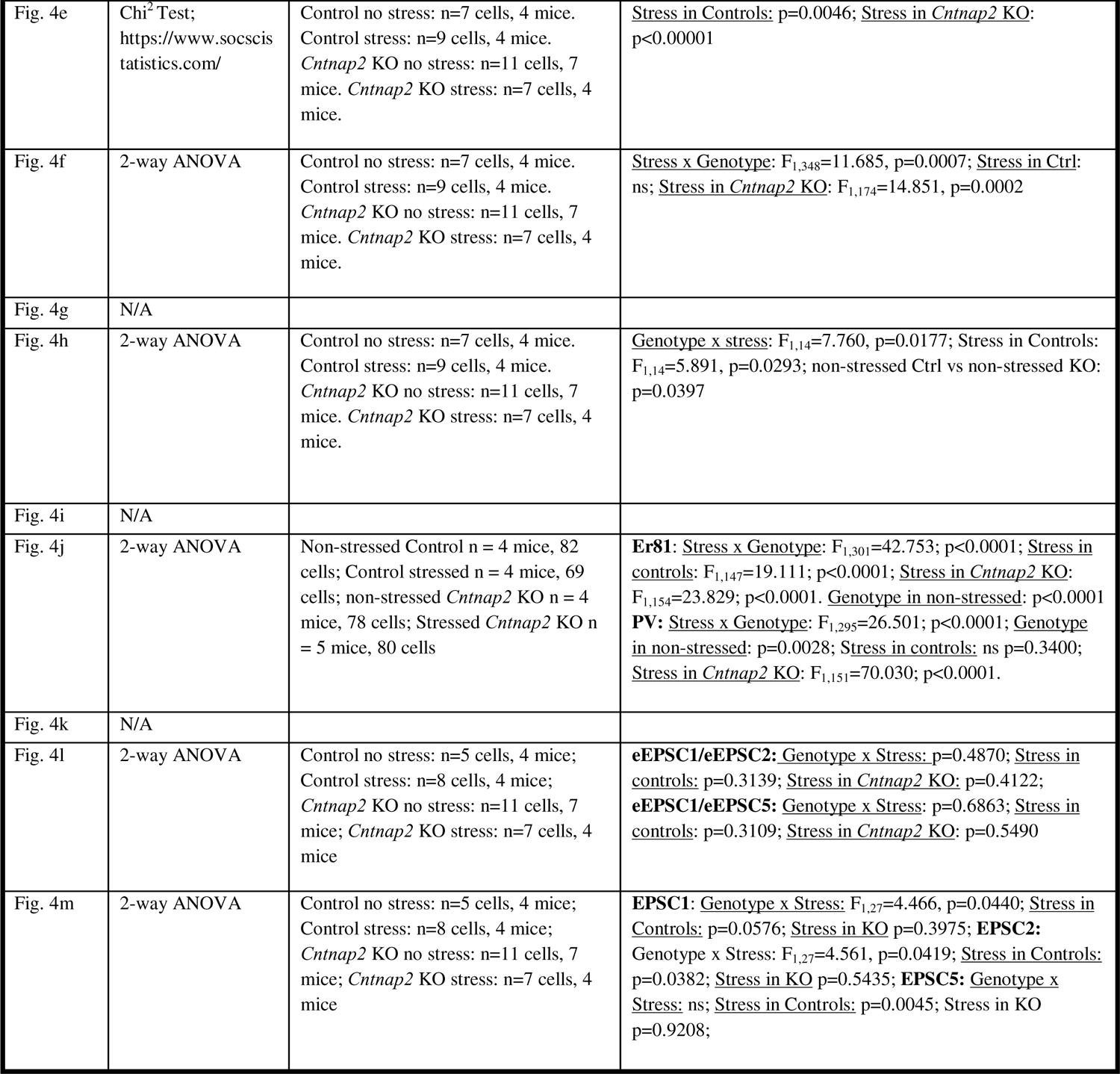

| Figure | Statistical Test | Sample size | p-Value |
| --- | --- | --- | --- |
| **Figure 1** |  |  |  |
| Fig. 1a | N/A |  |  |
| Fig. 1b | 2-way ANOVA | Non stressed Control: n=7; Stressed Control: n=11 | **Tone Test:** Genotype x Evolution of freezing interaction: $F_{2,32}=24.292$, $p<0.0001$; Inset (Tone Ratio): $F_{1,15}=24.418$, $p=0.0002$; **Context Test:** $F_{1,15}=4.721$, $p=0.0462$ |
| Fig. 1c | 2-way ANOVA | Non stressed *Cntnap2* KO: n=7; Stressed *Cntnap2* KO: n=7 | **Tone Test:** Genotype x Evolution of freezing interaction: $F_{2,24}=9.189$, $p=0.0011$; Inset (Tone Ratio): $F_{1,12}=35.235$, $p<0.0001$; **Context Test:** $F_{1,12}=5.116$, $p=0.0431$ |
| Fig. 1d | N/A |  |  |
| Fig. 1e | 2-way ANOVA | Control: n=18; *Cntnap2* KO: n=20 | **Tone test:** $p=0.9816$; Inset (Tone ratio): $p=0.1812$; **Context test:** $p=0.1442$ |
| Fig. 1f | 2-way ANOVA | Control: n=32; *Cntnap2* KO: n=34 | **Tone Test:** Genotype x Evolution of freezing interaction: $F_{2,128}=35.720$, $p<0.0001$; Inset (Tone ratio): $F_{1,64}=33.407$, $p<0.0001$; **Context Test:** $F_{1,64}=41.332$, $p<0.0001$ |
| Fig. 1g | 2-way ANOVA | Control *Pten*: n=7; *Pten* cKO: n=8<br>Control *Fmr1*: n=12; *Fmr1* KO: n=9<br>Control *Shank 3*: n=7; *Shank 3* KO: n=8 | **<u>Pten</u>: Tone Test:** Genotype x Evolution Freezing: $F_{1,26}=15.952$, $p<0.0001$; Inset (Tone Ratio): $F_{1,13}=22.571$, $p=0.0004$; **Context Test:** $F_{1,13}=6.498$, $p=0.0242$<br>**<u>Fmr1</u>: Tone Test:** Genotype x Evolution Freezing: $F_{2,38}=11.273$, $p<0.0001$; Inset (Tone Ratio): $F_{1,19}=41.062$, $p<0.0001$; **Context Test:** $F_{1,19}=38.148$, $p<0.0001$<br>**<u>Shank 3</u>: Tone Test:** Genotype x Evolution Freezing: $F_{2,26}=3.900$, $p=0.0330$; Inset (Tone Ratio): $F_{1,13}=5.955$, $p=0.0297$; **Context Test:** $F_{1,13}=9.073$, $p=0.0100$<br>**<u>All controls vs ASD models</u>:** Genotype x Evolution Freezing: $F_{6,94}=10.996$, $p<0.0001$; Inset (Tone Ratio): $F_{3,47}=19.475$, $p<0.0001$; **Context Test:** $F_{3,47}=17.941$, $p<0.0001$ |
| Fig. 1h | 2-way ANOVA | Control: n=11; *Cntnap2* KO: n=14 | Before: Control vs. KO $F_{1,23}=22.766$, $p=0.0002$; Before vs. after PTSD in KO: $F_{1,13}=36.965$, $p<0.0001$; Before vs. after PTSD in controls: ns; Genotype x Evolution interaction: $F_{1,23}=8.504$, $p=0.0078$; Mouse interaction Index: $p=0.0005$ |
| Fig. 1i | 2-way ANOVA; t-test | Control: n=11; *Cntnap2* KO: n=18 | Before: Control vs. KO $F_{1,27}=47.177$, $p<0.0001$; Before vs. After PTSD in Ctrl: ns; Before vs. After PTSD in KO: $F_{1,17}=106.643$, $p<0.0001$; Genotype x Evolution interaction: KO $F_{1,27}=31.494$, $p<0.0001$. RM Index: $p=0.0472$ |
| **Figure 2** |  |  |  |
| Fig. 2a | N/A |  |  |
| Fig. 2b | 2-way ANOVA | Control: n=19; *Cntnap2* KO: n=14 mice | <u>Genotype x Evolution interaction</u>: $F_{2,62}=60.528$, $p<0.0001$; Pre: $F_{1,31}=50.764$, $p<0.0001$; Inset: $F_{1,20}=71.695$, $p<0.0001$ |
| Fig. 2c | 2-way ANOVA | Control: n=19; *Cntnap2* KO: n=14 | **Tone test**: $p=0.7213$; inset: $p=0.4607$; **Context test**: $p=0.3777$ |
| Fig. 2d | 2-way ANOVA | Control: n=10; *Cntnap2* KO: n=12 | Post-ReC: Control vs. KO $F_{1,20}=6.808$, $p=0.0168$; Post-PTSD vs. Post-ReC in KO: $F_{1,11}=31.193$, $p=0.0002$, ns in Controls; <u>Genotype x Evolution interaction</u>: $F_{1,20}=9.622$, $p=0.0056$ |
| Fig. 2e | 2-way ANOVA | Control: n=10; *Cntnap2* KO: n=16 | Post-PTSD vs. Post-ReC in KO: $F_{1,15}=76.739$, $p<0.0001$, ns in Controls; <u>Genotype x Evolution interaction</u>: KO $F_{1,24}=59.508$, $p<0.0001$ |
| **Figure 3** |  |  |  |
| Fig. 3a | 2-way ANOVA | Control: n=7; *Cntnap2* KO: n=7 | mPFC: $F_{1,12}=5.469$, $p=0.0375$ |
| Fig. 3b | N/A |  |  |
| Fig. 3c | N/A |  |  |
| Fig. 3d | 2-way ANOVA | Control GFP: n=9; Control Chr2: n=7 | **Tone Test**: <u>Genotype x Evolution interaction</u>: $F_{2,28}=17.397$, $p<0.0001$; Inset: $F_{1,14}=35.729$, $p<0.0001$, **Context Test**: $F_{1,14}=9.162$, $p=0.009$ |
| Fig. 3e | 2-way ANOVA | Control GFP: n=9; *Cntnap2* KO GFP: n=11; *Cntnap2* KO ArchT: n=9 | **Tone Test**: <u>Genotype x Evolution interaction</u>: $F_{4,52}=7.523$, $p<0.0001$; Inset: $F_{2,26}=11.446$, $p=0.0003$, Control GFP vs. KO-GFP: $p<0.0001$; KO-GFP vs. KO-ArchT: $p=0.0364$. Pre-Tone, KO-GFP vs. ArchT: $p=0.1826$<br>**Context Test**: $F_{2,26}=4.352$, $p=0.0234$; Control GFP vs. KO-GFP: $p=0.0393$; KO-GFP vs. KO-ArchT: $p=0.011$ |
| Fig. 3f | 2-way ANOVA; t-test | *Cntnap2* KO PV-Cre GFP: n=9; *Cntnap2* KO PV-Cre Chr2: n=9 | **Tone Test**: <u>Genotype x Evolution interaction</u>: $F_{2,32}=33.793$, $p<0.0001$; Inset: $F_{1,16}=59.749$, $p<0.0001$, **Context Test**: $p=0.0005$ |
| Fig. 3g | 2-way ANOVA; t-test | *Cntnap2* KO PV-Cre GFP: n=9; *Cntnap2* KO PV-Cre Chr2: n=9 | **Tone Test**: <u>Genotype x Evolution interaction</u>: $F_{2,32}=33.793$, $p<0.0001$; Inset: $F_{1,16}=59.749$, $p<0.0001$, **Context Test**: $p=0.0005$ |
| Fig. 4a | N/A |  |  |
| Fig. 4b | 2-way ANOVA | Control no stress: n=7 cells, 4 mice. Control stress: n=9 cells, 4 mice. *Cntnap2* KO no stress: n=11 cells, 7 mice. *Cntnap2* KO stress: n=7 cells, 4 mice. | <u>Evolution of spike rate x Stress</u>: $F_{1,11}=31.193$, $p=0.0002$. In *Cntnap2* KO <u>Evolution of spike rate x Stress</u>: $F_{30,480}=1.560$, $p=0.0315$; in Controls, <u>Evolution of spike rate x Stress</u>: $F_{30,480}=1.560$, $p=0.5014$, |
| Fig. 4c | N/A |  |  |
| Fig. 4d | 2-way ANOVA; t-test | Control no stress: n=7 cells, 4 mice. Control stress: n=13 cells, 4 mice. *Cntnap2* KO no stress: n=12 cells, 7 mice. *Cntnap2* KO stress: n=8 cells, 4 mice. | <u>Stress x Genotype</u>: $F_{1,36}=9.269$, $p=0.0043$; Stress in Control: $p=0.2230$; Stress in *Cntnap2* KO: $p=0.0023$ |
| Fig. 4e | Chi<sup>2</sup> Test;<br>https://www.socscistatistics.com/ | Control no stress: n=7 cells, 4 mice.<br>Control stress: n=9 cells, 4 mice.<br>*Cntnap2* KO no stress: n=11 cells, 7 mice. *Cntnap2* KO stress: n=7 cells, 4 mice. | <u>Stress in Controls</u>: p=0.0046; <u>Stress in *Cntnap2* KO</u>: p<0.00001 |
| Fig. 4f | 2-way ANOVA | Control no stress: n=7 cells, 4 mice.<br>Control stress: n=9 cells, 4 mice.<br>*Cntnap2* KO no stress: n=11 cells, 7 mice. *Cntnap2* KO stress: n=7 cells, 4 mice. | <u>Stress x Genotype</u>: F<sub>1,348</sub>=11.685, p=0.0007; <u>Stress in Ctrl</u>: ns; <u>Stress in *Cntnap2* KO</u>: F<sub>1,174</sub>=14.851, p=0.0002 |
| Fig. 4g | N/A |  |  |
| Fig. 4h | 2-way ANOVA | Control no stress: n=7 cells, 4 mice.<br>Control stress: n=9 cells, 4 mice.<br>*Cntnap2* KO no stress: n=11 cells, 7 mice. *Cntnap2* KO stress: n=7 cells, 4 mice. | <u>Genotype x stress</u>: F<sub>1,14</sub>=7.760, p=0.0177; <u>Stress in Controls</u>: F<sub>1,14</sub>=5.891, p=0.0293; non-stressed Ctrl vs non-stressed KO: p=0.0397 |
| Fig. 4i | N/A |  |  |
| Fig. 4j | 2-way ANOVA | Non-stressed Control n = 4 mice, 82 cells; Control stressed n = 4 mice, 69 cells; non-stressed *Cntnap2* KO n = 4 mice, 78 cells; Stressed *Cntnap2* KO n = 5 mice, 80 cells | **Er81**: <u>Stress x Genotype</u>: F<sub>1,301</sub>=42.753; p<0.0001; <u>Stress in controls</u>: F<sub>1,147</sub>=19.111; p<0.0001; <u>Stress in *Cntnap2* KO</u>: F<sub>1,154</sub>=23.829; p<0.0001. <u>Genotype in non-stressed</u>: p<0.0001<br>**PV**: <u>Stress x Genotype</u>: F<sub>1,295</sub>=26.501; p<0.0001; <u>Genotype in non-stressed</u>: p=0.0028; <u>Stress in controls</u>: ns p=0.3400; <u>Stress in *Cntnap2* KO</u>: F<sub>1,151</sub>=70.030; p<0.0001. |
| Fig. 4k | N/A |  |  |
| Fig. 4l | 2-way ANOVA | Control no stress: n=5 cells, 4 mice;<br>Control stress: n=8 cells, 4 mice;<br>*Cntnap2* KO no stress: n=11 cells, 7 mice; *Cntnap2* KO stress: n=7 cells, 4 mice | **eEPSC1/eEPSC2**: <u>Genotype x Stress</u>: p=0.4870; <u>Stress in controls</u>: p=0.3139; <u>Stress in *Cntnap2* KO</u>: p=0.4122;<br>**eEPSC1/eEPSC5**: <u>Genotype x Stress</u>: p=0.6863; <u>Stress in controls</u>: p=0.3109; <u>Stress in *Cntnap2* KO</u>: p=0.5490 |
| Fig. 4m | 2-way ANOVA | Control no stress: n=5 cells, 4 mice;<br>Control stress: n=8 cells, 4 mice;<br>*Cntnap2* KO no stress: n=11 cells, 7 mice; *Cntnap2* KO stress: n=7 cells, 4 mice | **EPSC1**: <u>Genotype x Stress</u>: F<sub>1,27</sub>=4.466, p=0.0440; <u>Stress in Controls</u>: p=0.0576; <u>Stress in KO</u> p=0.3975; **EPSC2**: <u>Genotype x Stress</u>: F<sub>1,27</sub>=4.561, p=0.0419; <u>Stress in Controls</u>: p=0.0382; <u>Stress in KO</u> p=0.5435; **EPSC5**: <u>Genotype x Stress</u>: ns; <u>Stress in Controls</u>: p=0.0045; <u>Stress in KO</u> p=0.9208; |

**Table 3.**
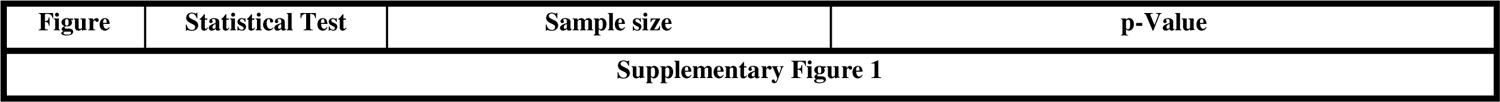
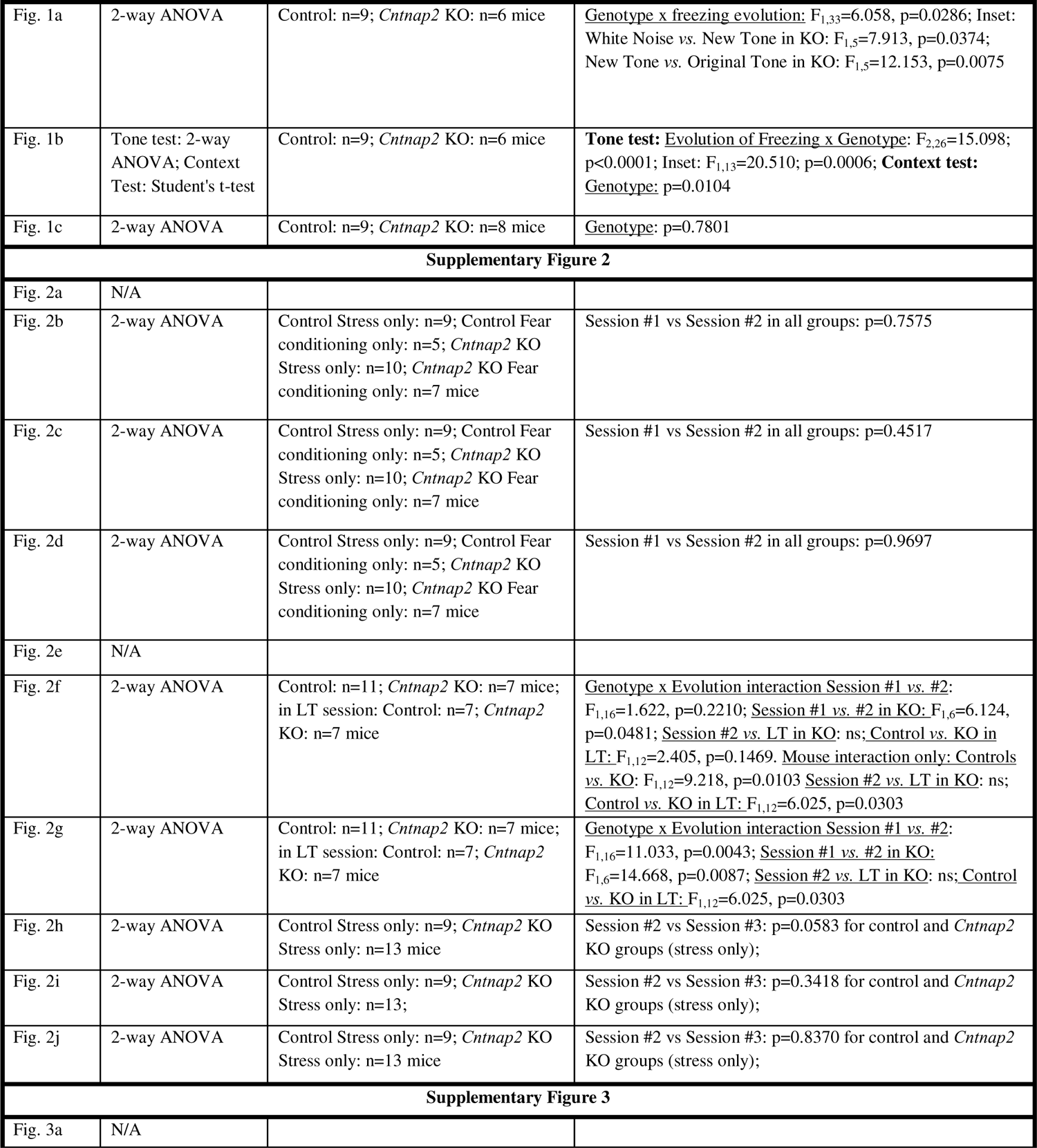
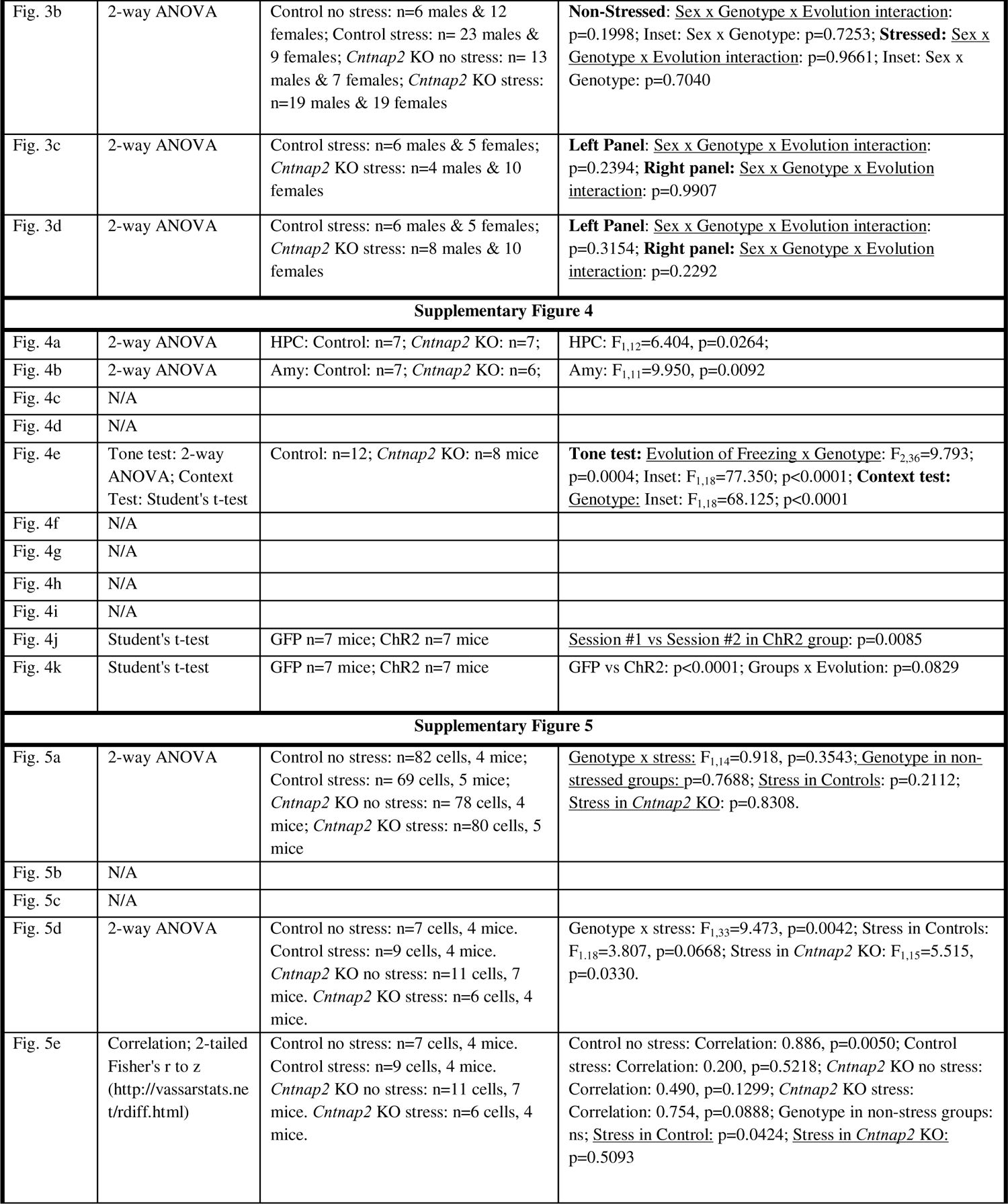
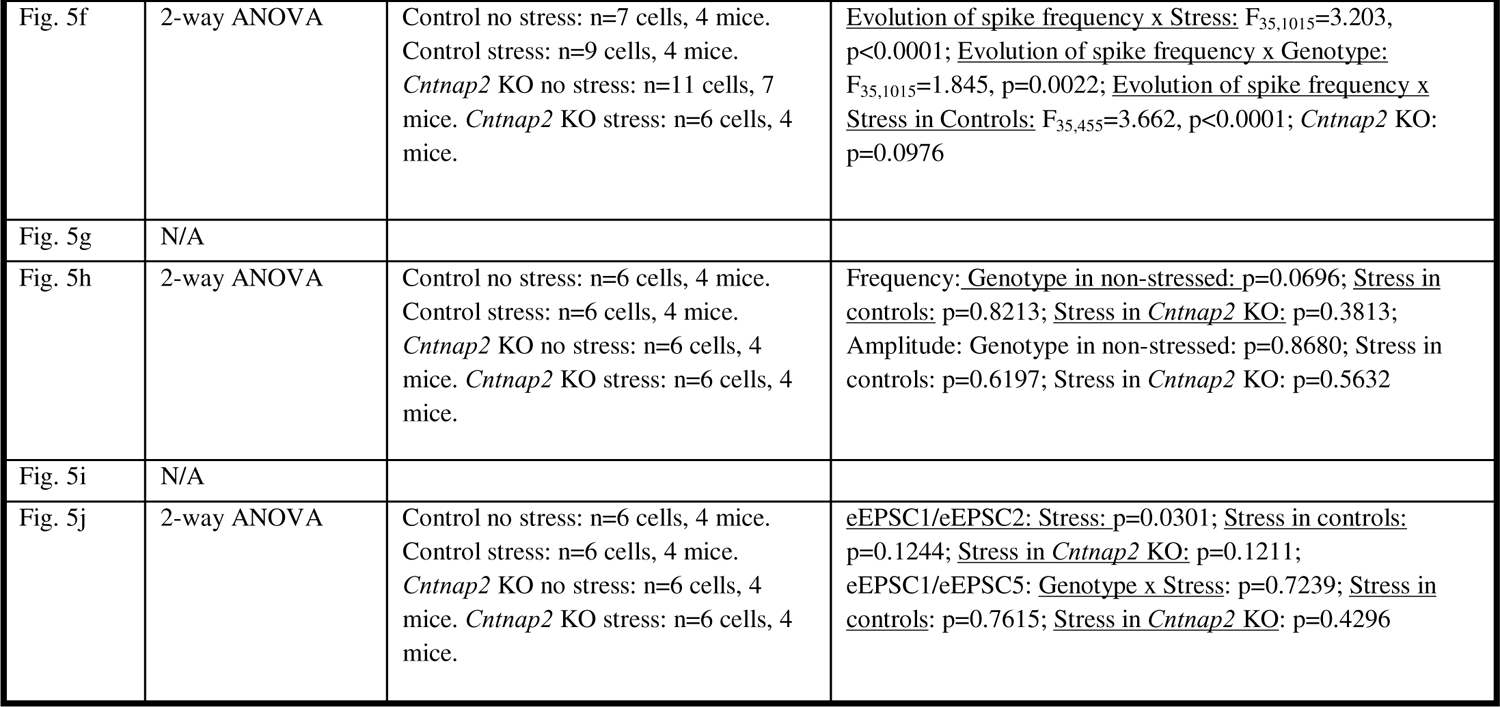

| Figure | Statistical Test | Sample size | p-Value |
| --- | --- | --- | --- |
| Supplementary Figure 1 |  |  |  |
| Fig. 1a | 2-way ANOVA | Control: n=9; *Cntnap2* KO: n=6 mice | <u>Genotype x freezing evolution</u>: $F_{1,33}=6.058$, $p=0.0286$; Inset: White Noise vs. New Tone in KO: $F_{1,5}=7.913$, $p=0.0374$; New Tone vs. Original Tone in KO: $F_{1,5}=12.153$, $p=0.0075$ |
| Fig. 1b | Tone test: 2-way ANOVA; Context Test: Student's t-test | Control: n=9; *Cntnap2* KO: n=6 mice | **Tone test**: <u>Evolution of Freezing x Genotype</u>: $F_{2,26}=15.098$; $p<0.0001$; Inset: $F_{1,13}=20.510$; $p=0.0006$; **Context test**: <u>Genotype</u>: $p=0.0104$ |
| Fig. 1c | 2-way ANOVA | Control: n=9; *Cntnap2* KO: n=8 mice | <u>Genotype</u>: $p=0.7801$ |
| Supplementary Figure 2 |  |  |  |
| Fig. 2a | N/A |  |  |
| Fig. 2b | 2-way ANOVA | Control Stress only: n=9; Control Fear conditioning only: n=5; *Cntnap2* KO Stress only: n=10; *Cntnap2* KO Fear conditioning only: n=7 mice | Session #1 vs Session #2 in all groups: $p=0.7575$ |
| Fig. 2c | 2-way ANOVA | Control Stress only: n=9; Control Fear conditioning only: n=5; *Cntnap2* KO Stress only: n=10; *Cntnap2* KO Fear conditioning only: n=7 mice | Session #1 vs Session #2 in all groups: $p=0.4517$ |
| Fig. 2d | 2-way ANOVA | Control Stress only: n=9; Control Fear conditioning only: n=5; *Cntnap2* KO Stress only: n=10; *Cntnap2* KO Fear conditioning only: n=7 mice | Session #1 vs Session #2 in all groups: $p=0.9697$ |
| Fig. 2e | N/A |  |  |
| Fig. 2f | 2-way ANOVA | Control: n=11; *Cntnap2* KO: n=7 mice; in LT session: Control: n=7; *Cntnap2* KO: n=7 mice | <u>Genotype x Evolution interaction Session #1 vs. #2</u>: $F_{1,16}=1.622$, $p=0.2210$; <u>Session #1 vs. #2 in KO</u>: $F_{1,6}=6.124$, $p=0.0481$; <u>Session #2 vs. LT in KO</u>: ns; <u>Control vs. KO in LT</u>: $F_{1,12}=2.405$, $p=0.1469$. <u>Mouse interaction only: Controls vs. KO</u>: $F_{1,12}=9.218$, $p=0.0103$ <u>Session #2 vs. LT in KO</u>: ns; <u>Control vs. KO in LT</u>: $F_{1,12}=6.025$, $p=0.0303$ |
| Fig. 2g | 2-way ANOVA | Control: n=11; *Cntnap2* KO: n=7 mice; in LT session: Control: n=7; *Cntnap2* KO: n=7 mice | <u>Genotype x Evolution interaction Session #1 vs. #2</u>: $F_{1,16}=11.033$, $p=0.0043$; <u>Session #1 vs. #2 in KO</u>: $F_{1,6}=14.668$, $p=0.0087$; <u>Session #2 vs. LT in KO</u>: ns; <u>Control vs. KO in LT</u>: $F_{1,12}=6.025$, $p=0.0303$ |
| Fig. 2h | 2-way ANOVA | Control Stress only: n=9; *Cntnap2* KO Stress only: n=13 mice | Session #2 vs Session #3: $p=0.0583$ for control and *Cntnap2* KO groups (stress only); |
| Fig. 2i | 2-way ANOVA | Control Stress only: n=9; *Cntnap2* KO Stress only: n=13; | Session #2 vs Session #3: $p=0.3418$ for control and *Cntnap2* KO groups (stress only); |
| Fig. 2j | 2-way ANOVA | Control Stress only: n=9; *Cntnap2* KO Stress only: n=13 mice | Session #2 vs Session #3: $p=0.8370$ for control and *Cntnap2* KO groups (stress only); |
| Supplementary Figure 3 |  |  |  |
| Fig. 3a | N/A |  |  |
| Fig. 3b | 2-way ANOVA | Control no stress: n=6 males & 12 females; Control stress: n= 23 males & 9 females; *Cntnap2* KO no stress: n= 13 males & 7 females; *Cntnap2* KO stress: n=19 males & 19 females | **Non-Stressed:** <u>Sex x Genotype x Evolution interaction:</u> p=0.1998; Inset: Sex x Genotype: p=0.7253; **Stressed:** <u>Sex x Genotype x Evolution interaction:</u> p=0.9661; Inset: Sex x Genotype: p=0.7040 |
| Fig. 3c | 2-way ANOVA | Control stress: n=6 males & 5 females; *Cntnap2* KO stress: n=4 males & 10 females | **Left Panel:** <u>Sex x Genotype x Evolution interaction:</u> p=0.2394; **Right panel:** <u>Sex x Genotype x Evolution interaction:</u> p=0.9907 |
| Fig. 3d | 2-way ANOVA | Control stress: n=6 males & 5 females; *Cntnap2* KO stress: n=8 males & 10 females | **Left Panel:** <u>Sex x Genotype x Evolution interaction:</u> p=0.3154; **Right panel:** <u>Sex x Genotype x Evolution interaction:</u> p=0.2292 |
| Supplementary Figure 4 |  |  |  |
| Fig. 4a | 2-way ANOVA | HPC: Control: n=7; *Cntnap2* KO: n=7; | HPC: $F_{1,12}=6.404$, p=0.0264; |
| Fig. 4b | 2-way ANOVA | Amy: Control: n=7; *Cntnap2* KO: n=6; | Amy: $F_{1,11}=9.950$, p=0.0092 |
| Fig. 4c | N/A |  |  |
| Fig. 4d | N/A |  |  |
| Fig. 4e | Tone test: 2-way ANOVA; Context Test: Student's t-test | Control: n=12; *Cntnap2* KO: n=8 mice | **Tone test:** Evolution of Freezing x Genotype: $F_{2,36}=9.793$; p=0.0004; Inset: $F_{1,18}=77.350$; p<0.0001; **Context test:** <u>Genotype:</u> Inset: $F_{1,18}=68.125$; p<0.0001 |
| Fig. 4f | N/A |  |  |
| Fig. 4g | N/A |  |  |
| Fig. 4h | N/A |  |  |
| Fig. 4i | N/A |  |  |
| Fig. 4j | Student's t-test | GFP n=7 mice; Chr2 n=7 mice | Session #1 vs Session #2 in Chr2 group: p=0.0085 |
| Fig. 4k | Student's t-test | GFP n=7 mice; Chr2 n=7 mice | GFP vs Chr2: p<0.0001; Groups x Evolution: p=0.0829 |
| Supplementary Figure 5 |  |  |  |
| Fig. 5a | 2-way ANOVA | Control no stress: n=82 cells, 4 mice; Control stress: n= 69 cells, 5 mice; *Cntnap2* KO no stress: n= 78 cells, 4 mice; *Cntnap2* KO stress: n=80 cells, 5 mice | <u>Genotype x stress:</u> $F_{1,14}=0.918$, p=0.3543; <u>Genotype in non-stressed groups:</u> p=0.7688; <u>Stress in Controls:</u> p=0.2112; <u>Stress in *Cntnap2* KO:</u> p=0.8308. |
| Fig. 5b | N/A |  |  |
| Fig. 5c | N/A |  |  |
| Fig. 5d | 2-way ANOVA | Control no stress: n=7 cells, 4 mice. Control stress: n=9 cells, 4 mice. *Cntnap2* KO no stress: n=11 cells, 7 mice. *Cntnap2* KO stress: n=6 cells, 4 mice. | Genotype x stress: $F_{1,33}=9.473$, p=0.0042; Stress in Controls: $F_{1,18}=3.807$, p=0.0668; Stress in *Cntnap2* KO: $F_{1,15}=5.515$, p=0.0330. |
| Fig. 5e | Correlation; 2-tailed Fisher's r to z (http://vassarstats.net/rdiff.html) | Control no stress: n=7 cells, 4 mice. Control stress: n=9 cells, 4 mice. *Cntnap2* KO no stress: n=11 cells, 7 mice. *Cntnap2* KO stress: n=6 cells, 4 mice. | Control no stress: Correlation: 0.886, p=0.0050; Control stress: Correlation: 0.200, p=0.5218; *Cntnap2* KO no stress: Correlation: 0.490, p=0.1299; *Cntnap2* KO stress: Correlation: 0.754, p=0.0888; Genotype in non-stress groups: ns; <u>Stress in Control:</u> p=0.0424; <u>Stress in *Cntnap2* KO:</u> p=0.5093 |
| Fig. 5f | 2-way ANOVA | Control no stress: n=7 cells, 4 mice.<br>Control stress: n=9 cells, 4 mice.<br>*Cntnap2* KO no stress: n=11 cells, 7 mice. *Cntnap2* KO stress: n=6 cells, 4 mice. | <u>Evolution of spike frequency x Stress</u>: $F_{35,1015}=3.203$, $p<0.0001$; <u>Evolution of spike frequency x Genotype</u>: $F_{35,1015}=1.845$, $p=0.0022$; <u>Evolution of spike frequency x Stress in Controls</u>: $F_{35,455}=3.662$, $p<0.0001$; *Cntnap2* KO: $p=0.0976$ |
| Fig. 5g | N/A |  |  |
| Fig. 5h | 2-way ANOVA | Control no stress: n=6 cells, 4 mice.<br>Control stress: n=6 cells, 4 mice.<br>*Cntnap2* KO no stress: n=6 cells, 4 mice. *Cntnap2* KO stress: n=6 cells, 4 mice. | Frequency: <u>Genotype in non-stressed</u>: $p=0.0696$; <u>Stress in controls</u>: $p=0.8213$; <u>Stress in *Cntnap2* KO</u>: $p=0.3813$; Amplitude: Genotype in non-stressed: $p=0.8680$; Stress in controls: $p=0.6197$; Stress in *Cntnap2* KO: $p=0.5632$ |
| Fig. 5i | N/A |  |  |
| Fig. 5j | 2-way ANOVA | Control no stress: n=6 cells, 4 mice.<br>Control stress: n=6 cells, 4 mice.<br>*Cntnap2* KO no stress: n=6 cells, 4 mice. *Cntnap2* KO stress: n=6 cells, 4 mice. | <u>eEPSC1/eEPSC2: Stress</u>: $p=0.0301$; <u>Stress in controls</u>: $p=0.1244$; <u>Stress in *Cntnap2* KO</u>: $p=0.1211$; <u>eEPSC1/eEPSC5: Genotype x Stress</u>: $p=0.7239$; <u>Stress in controls</u>: $p=0.7615$; <u>Stress in *Cntnap2* KO</u>: $p=0.4296$ |

## Data availability

All data are available in the main text or the supplementary materials. Raw data will be made available upon reasonable request.

## DISCUSSION

Overall, this work uncovers a reciprocal relationship between PTSD-like memory formation and the core traits of ASD. Whilst deficits in stress processing are well-known in ASD, there has been a lack of research into the risk of PTSD development in autism, that stems from a poor understanding of the combined disorders and the absence of suitable detection tools ^9,39,40^. Here, we demonstrate in four mouse models of ASD that, contrary to a control population in which PTSD is triggered by an *extreme* stress ^1^, a single *mild* stress in ASD is sufficient to produce traumatic memory. This study therefore provides the first direct demonstration that ASD is associated with a risk of developing PTSD-like memory for mildly stressful situations and that interneurons are key cellular substrates to traumatic memory. Because ASD mice models displayed intact contextual memory capacities in non-stressful situations, we suggest that an impaired ability to cope with stressful situations is responsible for the switch from normal to PTSD-like memory. One of the limitations to consider with this new animal model is that qualitative aspects of PTSD (e.g., anxiety, avoidance of reminders of the event, emotional numbing) were not analyzed. Future studies should aim to further dissect these important aspects that are relevant to human behavior.

Human functional imaging and animal model studies suggest that cortical circuit abnormalities in ASD contribute to altered sensory representations and difficulties in stress coping. Our analysis uncovers the specific circuit alterations underlying the hypersensitivity to stress that leads to PTSD-like memory formation in ASD and allows the reinterpretation of the role of the circuitry responsible for PTSD-like memory formation in general. PTSD is indeed thought to be driven by amygdala hyperfunctioning, underlying the observed excessive fear. Yet, we observed the development of PTSD-like memory, associated with <u>hypo</u>activation of the amygdala in the *Cntnap2 KO* mice. Interestingly, we also found hippocampus hypoactivation, likely underlying contextual amnesia ^22^. Together, our results confirm the involvement of the hippocampus and contextualization in determining the nature of the memory formed during a stressful event in both ASD and PTSD. This work also calls for reevaluating the mechanisms underlying traumatic memory formation, which we demonstrate can be triggered by mPFC hyperactivation during trauma, combined with an acute mild stress in ASD mouse models (**Fig. 1 and 3**), and not by mPFC hypoactivation, as currently established for PTSD (Kredlow et al., 2022; Elzinga and Bremner, 2002). We therefore provide new ground to further dissect PTSD and its overlap with ASD pathophysiology.

Behaviorally, ASD and PTSD display similar characteristics ^26^, including impaired emotional regulation, cognitive rigidity, and fragmented autobiographical memory ^5^. Anatomically, the mPFC, crucial for executive functions, has been implicated in the pathophysiology of both ASD ^13,18^ and PTSD (Kredlow et al., 2021; Dahlgren et al., 2018). We show here that traumatic memory in ASD aggravates social impairments and repetitive behaviors, likely due to dysregulated cortico-striatal circuits top-down control from the mPFC ^43^. We reveal how cell input-output function is modulated in ASD, whereby the homeostatic capabilities of cortical interneurons are altered after stress, subsequently showing that the PV-INs of the mPFC are directly responsible for traumatic memory formation in ASD. In response to stress, PV-INs in the control condition are more excitable (*i.e.* decreased latency to first spike, increased firing frequency; **Fig.4**), increasing their inhibitory power. This is a necessary adaptive response, mediating the fine-tuned activation of the mPFC during stress (Jacobs and Moghaddam, 2021).However, PV-INs in the *Cntnap2* KO condition do not display this adaptive response, instead showing decreased excitability following stress (*i.e.* increased latency to first spike, decreased firing frequency; **Fig.4**), and thus a lower inhibitory power within the circuit ^33,45^. These deficits therefore underlie mPFC hyperactivation, preventing the *Cntnap2* KO mice from adapting adequately to stress, and in turn leading to PTSD-like memory formation (**Graphical abstract**). It would be of interest to further describe the role of PV-INs in PTSD, as this investigation has yet to be undertaken, and could uncover additional convergent mechanisms for ASD and PTSD centering around this interneuron population. In addition, PV-INs pose as potential targets for therapeutic intervention as they have been shown to restore balance in the mPFC circuit ^18^. Modulating the activity of the PV-INs in mice models of ASD would be an insightful investigation to restore normal memory and reduce the core symptoms of ASD.

The therapeutic approach explored here, recontextualization, reveals that pathological memory can be reshaped into adapted fear memory. By reactivating the traumatic memory in the original environment, recontextualization allows the re-allocation of trauma representation into specific context, thereby suppressing abnormal hypermnesia ^22^. In a clinical perspective though, our results suggest that recontextualization could cure PTSD-related memory in ASD. While this is promising, one caveat for the implementation of this procedure in patients will arise from being able to identify the origin of the trauma and detect PTSD in ASD. Our results indeed support that everyday life situations could be experienced as traumatic in ASD patients, as previously suggested from clinical studies ^8,46^. As new methods of detection emerge ^47^, our study calls for better use of predictive tools that enable efficient risk assessment and early interventions of PTSD among those likely to experience trauma, such as patients with autism. Timely detection appears essential since PTSD worsens the core symptoms of ASD and is strongly associated with various psychiatric comorbidities and suicide ^48^.

Overall, our study provides a new tool to further dissect the overlap between PTSD and ASD and the underlying alterations in emotional memory. It encourages future research to enhance detection and implementation of therapeutic strategies to pave the way towards alleviating the uncontrollable reactivation of traumatic memory in autism in the context of environmental pressure such as the COVID-19 pandemic ^49^, lockdowns and curfews which disrupt routines and may produce long-lasting cognitive impairments in vulnerable populations.

## Abbreviations

ASD: Autism Spectrum Disorder

KO: Knock-Out

mPFC: Medial PreFrontal Cortex

PTSD: Post-Traumatic Stress Disorder

PV-INs: Parvalbumin Interneurons

## Acknowledgments

We would like to thank Dr Andreas Frick for the *Pten* cKO and *Fmr1* KO mice and Drs Nathalie Sans and Mireille Montcouquiol for the *Shank3* KO. We thank all the personnel of The Australian Phenomics Facility and of The Neurocentre Magendie involved in mouse care.

## Author contributions

Conceptualization: ND, ASA; Methodology: ND, ASA, AD; Investigation: ASA, TVA, YS, AS, NYA Supervision: ND, ASA; Writing—original draft: ND, ASA; Writing—review & editing: ND, ASA, NYA, AM, AD

## Supplementary Materials

Fig. S1 to S5

Graphical Abstract

Table S1 to S3

