## Supplementary Figures for "Parvalbumin Interneuron Activity Underlies Vulnerability to Post-Traumatic Stress Disorder in Autism"

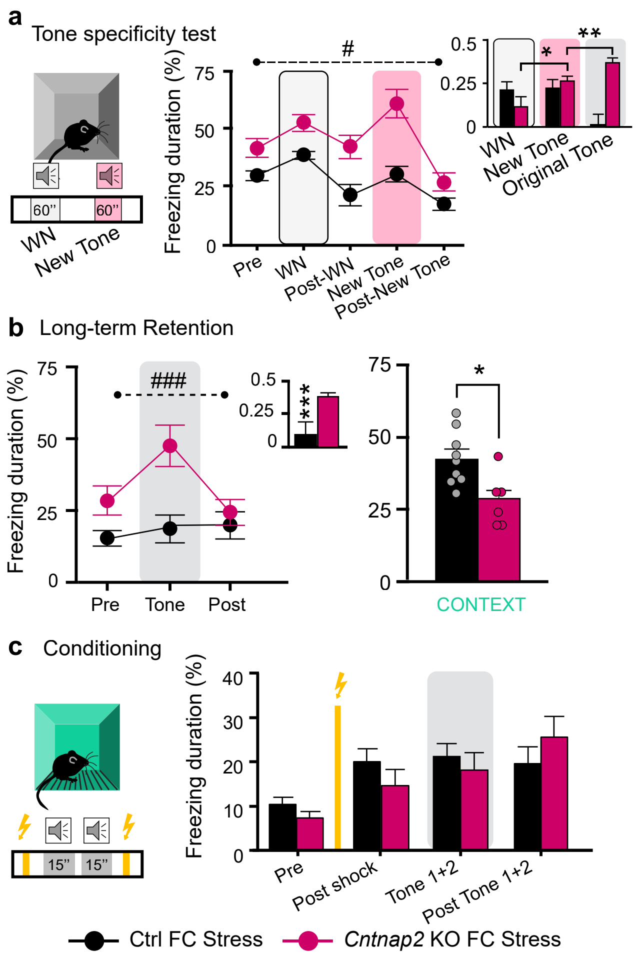


Supplementary Figure 1

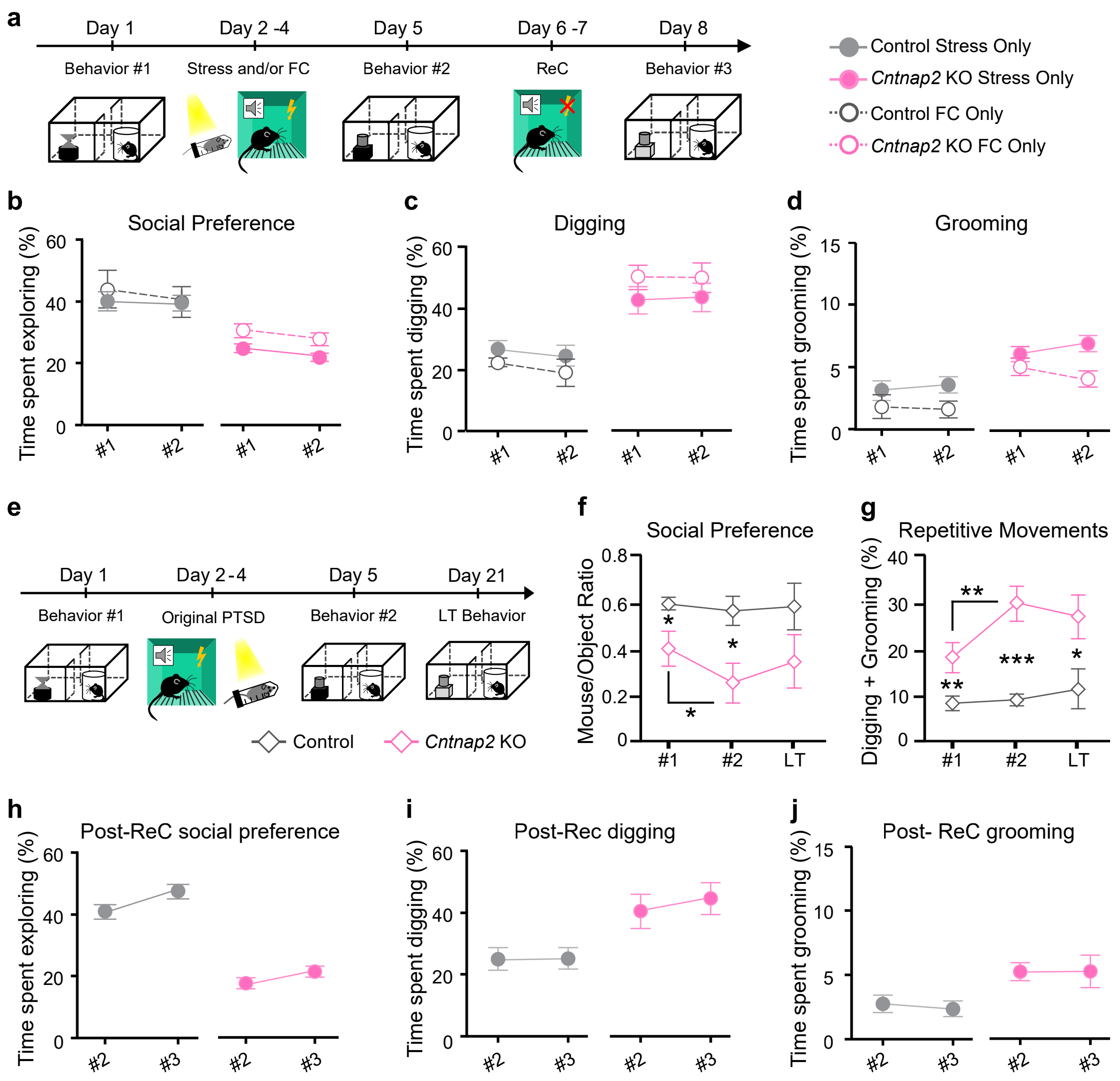


### Supplementary Figure 2

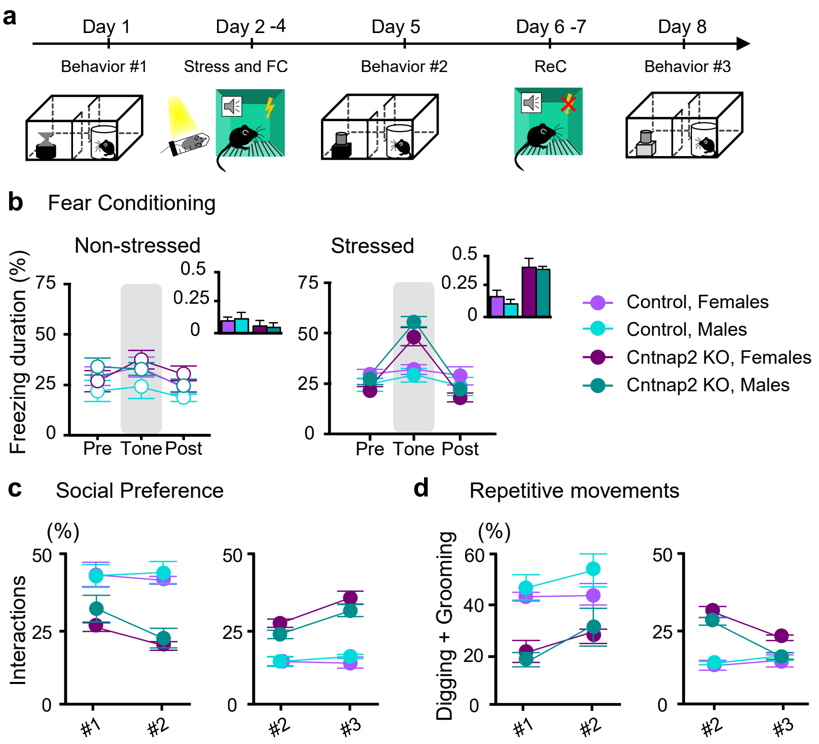


Supplementary Figure 3

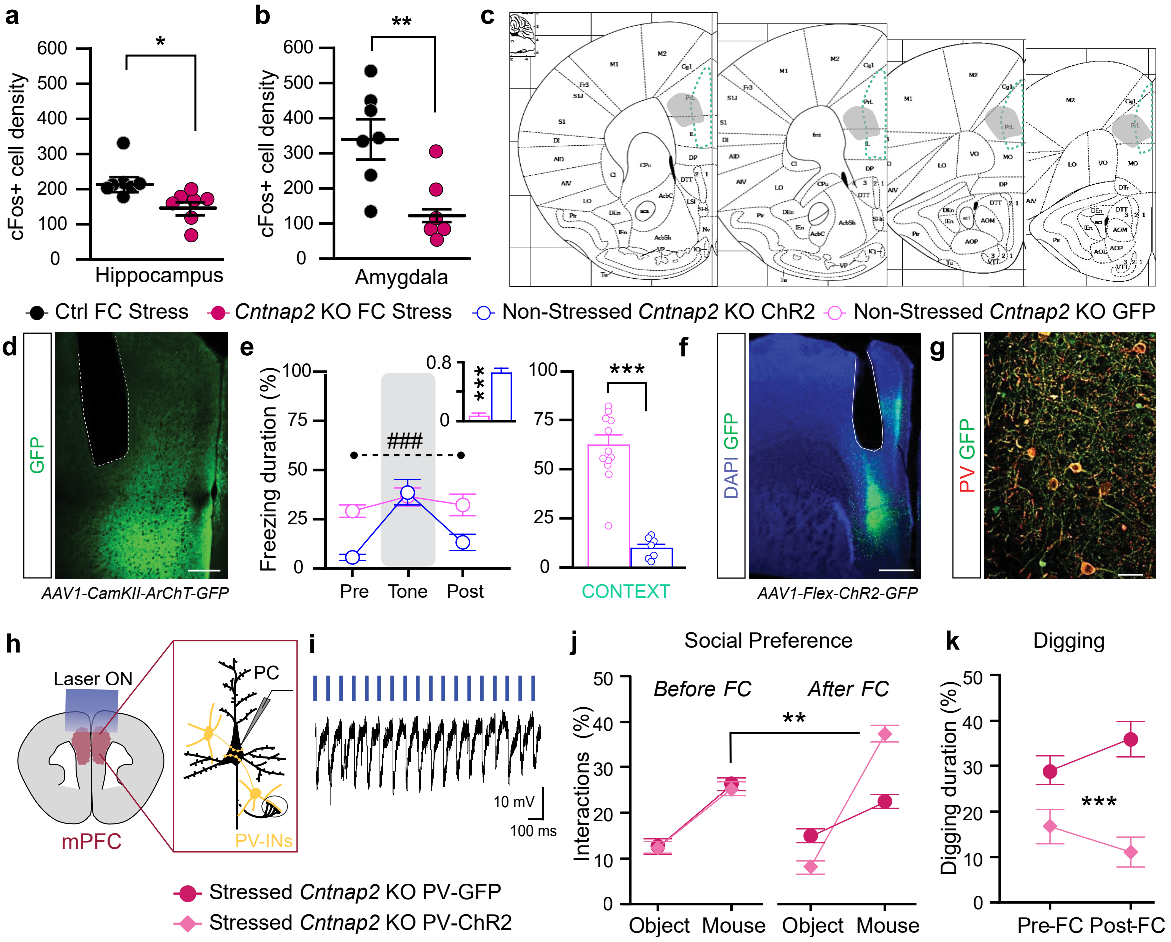


Supplementary Figure 4

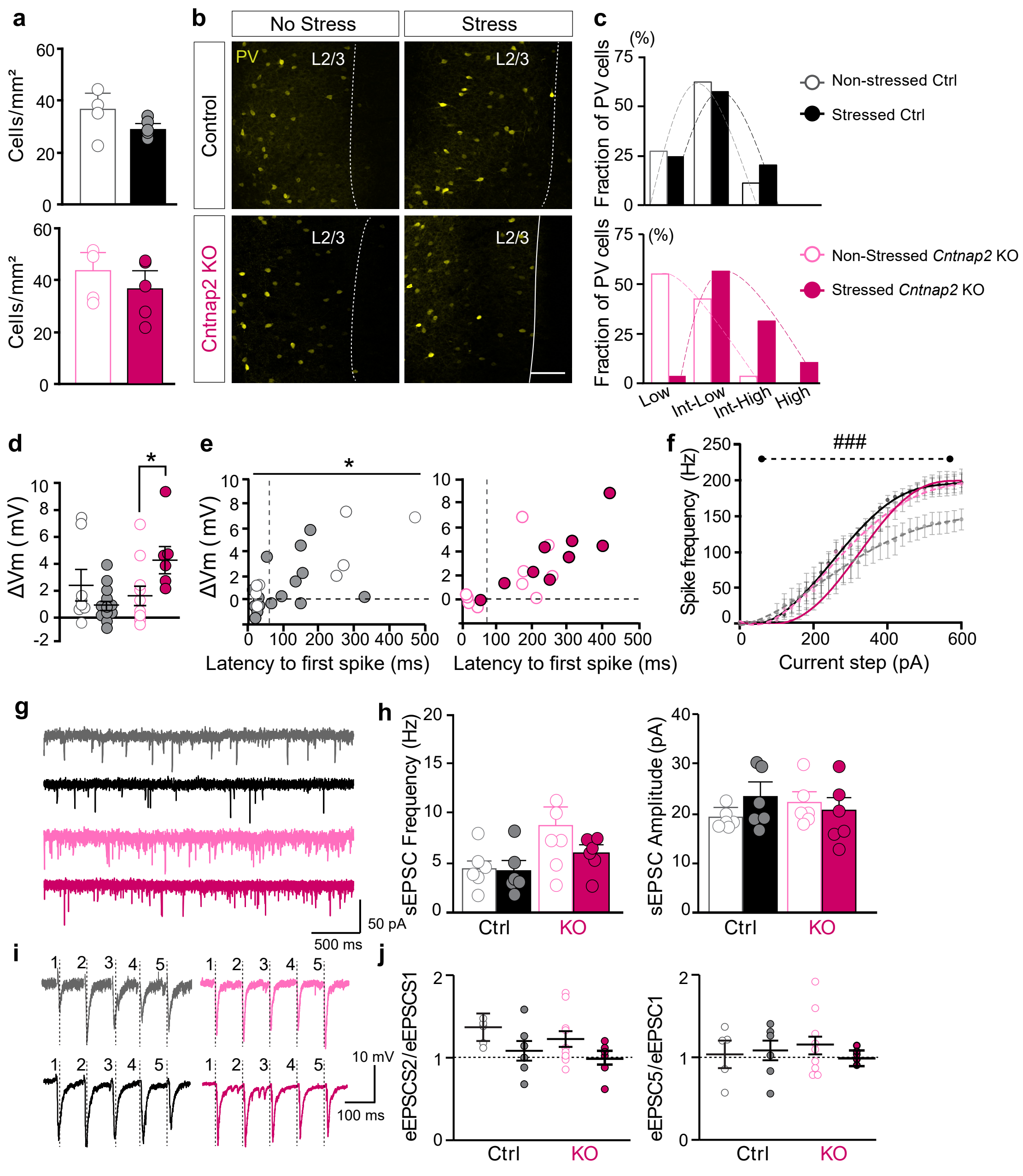


Supplementary Figure 5

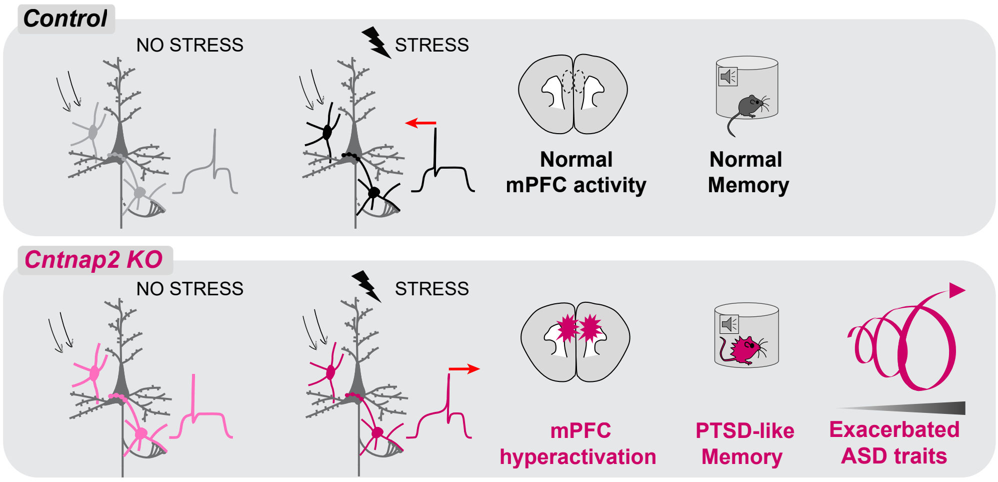


Graphical Abstract
