## Supplementary Legends for "Parvalbumin Interneuron Activity Underlies Vulnerability to Post-Traumatic Stress Disorder in Autism"

**Fig. S1. PTSD-like memory assessment.** **a,** Specificity of the fear response to the tone: mice were exposed to 2 tones: a white noise (WN), i.e., different from the conditioning tone, and a new tone (65dB, 3000Hz), i.e., similar to the conditioning tone (65dB, 1000Hz). Both controls and *Cntnap2* KO displayed low responses to the white noise. However, *Cntnap2* KO showed a stronger response to the new tone (#: Interaction white noise *vs.* new tone x genotype: <0.05), demonstrating the typical partial generalization observed in PTSD-like memory profile^1^. Inset: Tone discrimination ratio showing the fear response during each tone compared to freezing before and after each tone (white noise *vs.* new tone in KO: p<0.05; new tone *vs.* original tone in KO: p<0.01). **b,** 21 days after conditioning, *Cntnap2* KO mice (n=6) still displayed maladaptive memory (tone test: #: Interaction freezing x genotype: p<0.001; inset: genotype: p<0.001 context test: genotype: p<0.05). Control mice (n=9) demonstrated normal memory. **c,** Specific reactivity to the electric footshock: ns, no difference in freezing behavior between controls (n=18) and *Cntnap2* KO (n=20) mice throughout the conditioning session on Day1. Data are presented as mean ± SEM. ***: p<0.001; **: p<0.01; *: p<0.05 from 2-way ANOVA; see table for precise statistics.

**Fig. S2. Control behavioral experiments. a,** Timeline for behavioral testing during modified PTSD protocol presentation and recontextualization. **b,** Social preference experiment, representing the percentage of time spent exploring an unfamiliar mouse before (#1) and after (#2) PTSD protocol (*i.e*. stress + Fear Conditioning (FC)); control stress only: n=9; control FC only: n=5; *Cntnap2* KO stress only: n=10; *Cntnap2* KO FC only: n=7; ns for all)**. c,** Digging before and after PTSD protocol (control stress only: n=9; control FC only: n=5; *Cntnap2* KO stress only: n=10; *Cntnap2* KO FC only: n=7; ns for all). **d,** Grooming before and after PTSD protocol (control stress only: n=9; control FC only: n=5; *Cntnap2* KO stress only: n=10; *Cntnap2* KO FC only: n=7; ns for all). **e,** Timeline for behavioral testing for the long-term effects of the original PTSD paradigm. **f,** Social preference in control (n = 11) and *Cntnap2* KO (n = 7) mice across three behavioural sessions (#1 (before PTSD): p<0.05; #2 (after PTSD): p<0.05); LT (long term behavior): ns; *Cntnap2* KO #1 *vs.* #2: p<0.05, *Cntnap2* KO #2 *vs.* LT: ns, Mouse interaction only (not pictured): p<0.05). **g,** Repetitive movements in control and *Cntnap2* KO mice across three behavioral sessions (#1 (before PTSD): p<0.01; #2 (after PTSD): p<0.001); LT (long term behavior): p<0.05; *Cntnap2* KO #1 *vs.* #2: p<0.01). **f,** Social preference before (#2) and after recontextualization (Post-ReC, #3; control stress only: n=9; *Cntnap2* KO stress only: n=13; ns)**. g,** Digging before (#2) and after recontextualization (#3; control stress only: n=9; *Cntnap2* KO stress only: n=13; ns)**. h,** Grooming before (#2) and after recontextualization (#3; control stress only: n=9; *Cntnap2* KO Stress only: n=13; ns)**.** Data are presented as mean ± SEM. ***: p<0.001; **: p<0.01; *: p<0.05 from 2-way ANOVA. See table for precise statistics.

**Fig. S3. Sex differences analysis. a,** Timeline for behavioral testing during modified PTSD protocol presentation and recontextualization. FC: Fear Conditioning. ReC: recontextualization. **b,** Similar fear responses in males and females during the tone test, in non-stressed (left; ns; tone ratio (inset): ns) and stressed conditions (right; ns; tone ratio (inset): ns); control no stress: n=6 males & 12 females; control stress: n= 23 males & 9 females; *Cntnap2* KO no stress: n= 13 males & 7 females; *Cntnap2* KO stress: n=19 males & 19 females. **c**, Social preference test: duration of interaction with an unfamiliar mouse, in males and females before (#1) and after (#2) PTSD protocol (left), and before (#2) and after (#3) ReC protocol (right) (stress + FC; control stress: n=6 males & 5 females; *Cntnap2* KO stress: n=4 males & 10 females; ns). **d,** Repetitive movements in males and females before (#1) and after (#2) PTSD protocol (left), and before (#2) and after (#3) ReC protocol (right), (control stress: n=6 males & 5 females; *Cntnap2* KO stress: n=8 males & 10 females; ns). Data presented as mean ± SEM. ***: p<0.001; **: p<0.01; *: p<0.05 from 2-way ANOVA. See table for precise statistics.

**Fig. S4. mPFC photostimulation and controls experiments. a,b,** The density of cFos-expressing cells in the hippocampus **(a)** and amygdala **(b)** of stressed control (n = 7 mice) and stressed *Cntnap2* KO (n = 6 mice; hippocampus: p<0.05, amygdala: p<0.01). **c,** Schematics of viral infection areas (green dashed lines) from the posterior (Bregma +1.70 mm) to the anterior mPFC (Bregma +2.34 mm). The grey area represents the location of optic fiber implantation. **d,** Illustration of virus spread and probe placement for pyramidal cell photoinactivation (Fig 3e), 10x, scale: 250µm. **e,** Mean fear responses of non-stressed *Cntnap2* KO mice injected with either ChR2 (pyramidal cell stimulation; n = 8) or GFP (control; n = 12) during the tone test (left, p<0.001; Tone Ratio (inset): p<0.001) and context test (right, p<0.001). **f,** Illustration of virus spread and probe placement for PV-IN photostimulation (Fig. 3f; 4x, scale: 500µm). g**,** The colocalization between the AAV1-Flex-ChR2 virus (GFP, green) and the PV protein (red) in PV-Cre *Cntnap2* KO mice; 40x, scale: 50µm. **h,** Schematics of the *in vitro* electrophysiological recordings of the mPFC pyramidal neurons under PV-IN photostimulation (orange). **i,** Light-evoked post-synaptic inhibitory responses recorded in pyramidal cells in current clamp mode at -70 mV (n=3 cells). **j,** Social preference tests before and after PTSD protocol (FC) with PV-IN photostimulation during conditioning (Methods): duration of interaction with an unfamiliar mouse (n=7 GFP; n=7 ChR2; p<0.001). **k,** Time spent digging before and after modified PTSD protocol with PV-IN photostimulation during conditioning (Methods) (n=7 GFP; n=7 ChR2; Controls *vs.* KO: p<0.0001ns). Data are presented as mean ±SEM. ***: p<0.001; **: p<0.01; *: p<0.05 from 2-way ANOVA. See table for precise statistics.

**Fig. S5. PV-INs properties after stress. a,** PV-IN density in layer 2/3 of the mPFC in control (top; control no stress: n=4; control stress: n= 5; ns) and *Cntnap2* KO mice (bottom; *Cntnap2* KO no stress: n= 4; *Cntnap2* KO stress: n=5 mice; ns). **b,** Representative images of PV-IN density (yellow; scale: 100µm). **c,** Distribution of PV fluorescence intensities in PV-INs from control and *Cntnap2* KO mice (control no stress: n=4; control stress: n= 5; *Cntnap2* KO no stress: n=4; *Cntnap2* KO stress: n=5). **d,** Changes in membrane potential at near threshold, between the beginning and the end of the injected current step (∆Vm; see Methods). control no stress: n=7 cells; control stress: n=13 cells; *Cntnap2* KO no stress: n=11 cells; *Cntnap2* KO stress: n=6 cells. Stress in control: ns; stress in *Cntnap2* KO: p<0.05. **e**, Correlation between latency to first spike and change in membrane potential representing activation of the delayed rectifying current (∆Vm), showing an increase in non-delayed cells upon stress in control (left; n = 7 control no stress and 13 control stress; p<0.05) and a trend to decrease in non-delayed cells in the *Cntnap2* KO mice (right; n=11 KO no stress and 6 KO stress). Dashed grey line represents the limit between delayed *vs.* non-delayed cells (see Methods). **f,** Firing frequency as a function of current steps in PV-INs clamped at -70mV (control no stress: n=7 cells; control stress: n=13 cells; *Cntnap2* KO no stress: n=11 cells; *Cntnap2* KO stress: n=6 cells) is changed by stress in controls (p<0.001) and with genotype (p<0.01). **g,** Representative traces of PV-INs spontaneous Excitatory Post-Synaptic Currents (sEPSCs). **h,** Quantification of sEPSCs frequency (left) and amplitude (right), showing no difference in synaptic inputs onto PV-INs (control no stress: n=7 cells; control stress: n=13 cells; *Cntnap2* KO no stress: n=11 cells; *Cntnap2* KO stress: n=6 cells; ns; 100ms, 100mV). **i,** Representative traces of PV-IN evoked EPSCs (eEPSCs) in 20 Hz paired-pulse ratios (PPR); non-stressed (light grey), and stressed (black) controls, non-stressed (light pink), and stressed (dark pink) *Cntnap2* KO mice; scale 10ms/25pA). **j,** Quantification of PPR ratio for 1^st^ and 2^nd^ eEPSCs (left; ns) and 1^st^ and 5^th^ eEPSCs (right; ns); control no stress: n=7 cells; control stress: n=13 cells; *Cntnap2* KO no stress: n=11 cells; *Cntnap2* KO stress: n=6 cells. Data are presented as mean ± SEM. ***: p<0.001; **: p<0.01; *: p<0.05 from 2-way ANOVA, otherwise stated. See table for precise statistics.

**Graphical Abstract**. This study reveals a cellular mechanism underlying PTSD-like memory formation and concurrent worsening of the core symptoms in the *Cntnap2* KO mouse model of autism spectrum disorder (ASD). Top panel: In control mice, parvalbumin interneurons adjust their firing upon stress (red arrow) to maintain the excitation-inhibition balance in the mPFC circuitry, resulting in normal memory formation. Middle panel: In *Cntnap2* KO mice, PV-INs firing properties (light pink) resemble that of the stressed control condition (black, top panel). Exposure to stress further induces a significant shift (red arrow) in PV-IN firing properties in the *Cntnap2* KO condition (dark pink), which underlies mPFC hyperactivation. This likely disrupts the top-down control on downstream structures involved in the fear circuitry, overall altering memory processing, and triggering the formation of PTSD-like memory. Occurrence of traumatic memory correlates with the exacerbation of social impairments and repetitive behaviors (spiral).
